# Integration of QTL and transcriptome approaches for the identification of genes involved in tomato response to nitrogen deficiency

**DOI:** 10.1101/2023.10.26.564109

**Authors:** H. Desaint, A. Héreil, J. Belinchon-Moreno, Y. Carretero, E. Pelpoir, M. Pascal, M Brault, D. Dumont, F. Lecompte, P. Laugier, R. Duboscq, F. Bitton, M. Grumic, C. Giraud, P. Ferrante, G Giuliano, F. Sunseri, M. Causse

**Affiliations:** INRAE, UR1052 GAFL, 84143 Montfavet, France; INRAE, UR407, Pathologie Végétale, 84143 Montfavet, France; INRAE, UR115 PSH, 84914 Avignon, France; INRAE, UE A2M, 84143 Montfavet, France; Casaccia Research Centre, Italian National Agency for New Technologies, Energy, and Sustainable Economic Development, Rome, Italy; Mediterranean University of Reggio Calabria, Reggio Calabria, Italy

**Keywords:** *Solanum lycopersicum* L., abiotic stress, nitrogen, RNAseq analysis, quantitative trait loci (QTL), genome-wide association study (GWAS)

## Abstract

Optimising plant nitrogen (N) usage and inhibiting N leaching loss in the soil-crop system is crucial to maintain crop yield and reduce environmental pollution. This study aimed at identifying quantitative trait loci (QTL) and differential expressed genes (DEGs) between two N treatments in order to list candidate genes related to nitrogen-related contrasting traits in tomato varieties. We characterised a genetic diversity core-collection (CC) and a multi-parental advanced generation intercross (MAGIC) tomato population grown in greenhouse under two nitrogen levels and assessed several N-related traits and mapped QTLs. Transcriptome response under the two N conditions was also investigated through RNA sequencing of fruit and leaves in four parents of the MAGIC population.

Significant differences in response to N input reduction were observed at the phenotypic level for biomass and N-related traits. Twenty-seven (27) QTLs were detected for three target traits (Leaf N content, leaf Nitrogen Balance Index and petiole NO_3_^-^ content), ten and six at low and high N condition, respectively; while 19 QTLs were identified for plasticity traits. At the transcriptome level, 4,752 and 2,405 DEGs were detected between the two N conditions in leaves and fruits, respectively, among which 3,628 (50.6%) in leaves and 1,717 (71.4%) in fruit were genotype specific. When considering all the genotypes, 1,677 DEGs were shared between organs or tissues.

Finally, we integrated DEGs and QTLs analyses to identify the most promising candidate genes. The results highlighted a complex genetic architecture of N homeostasis in tomato and novel putative genes useful for breeding improved-NUE tomato.

**Highlight:** Tomato response to nitrogen deficiency is genetically controlled by a few QTLs and impacts the expression of a large number of genes, among which some are good targets for breeding sober varieties.

## Introduction

Nitrogen (N) is considered the most used fertilizer in cropping systems, accounting by weight for nearly 60% of all the applied fertilizers (FAOSTAT, 2022). Their use has been steadily increasing over the past 60 years, from a world usage of 11 Tg N year^−1^ in 1961 to 109 Tg N year^−1^ in 2021 (FAOSTAT, 2022). As a result, a dramatic increase in crop yield has been achieved, although accompanied by considerable negative impacts on the environment. Indeed, more than half of the N added to cropland is lost in the environment, leading to adverse environmental impacts including nitrate (NO_3_^-^) leaching, eutrophication of water surface as well as the emission of the greenhouse gas nitrous oxide (N_2_O) (Snyder et al., 2009). Compared to other farming systems, greenhouse vegetable production requires higher watering and N inputs. The absence of nutritive solution recirculation, resulted in massive NO_3_^-^ leaching loss and N_2_O emissions, estimated by a meta-analysis to be 64% and 137% higher than open-field vegetable production in China, respectively (Wang et al., 2018). Among the greenhouse-based vegetable production system, tomato is the most important crop in terms of cultivation area (FAOSTAT, 2022). It is also one of the most over-fertilized crops, with often a large difference between the optimal N rate and the actual N application rate (Ren et al., 2022).The European legislation to reduce NO_3_^-^ leachates led to many technical innovations to improve N management in the past decades. The most promising are closed-loop irrigation systems, which can reduce up to 75% the total N supply (Méndez-Cifuentes et al., 2020). However, apart from Netherlands, Belgium and France, the majority (> 90%) of soilless greenhouses in Europe are still free-draining (Incrocci et al. 2020). Indeed, aside from economic considerations, the main limitation of this system is the increasing salinization of the recirculation solution due to the saline irrigation water frequently occurring in many Mediterranean coastal areas (Magán et al., 2008). Beyond environmental issues related to the fertilizer overuse, the excessive N-fertilization implies other detrimental effects. Frequently, the N regime alters the tomato leaf metabolome and its relationship with pest response. As an example, high-N fertilization is linked to an increased attractiveness to *Bemisia tabaci* through an altered volatile compound emission (Islam et al., 2017). Furthermore, N over-fertilization does not significantly improve tomato yield, by contrast it reduces fruit quality by decreasing sugar and increasing acid content (Bénard et al., 2009; Truffault et al., 2019). All these considerations support the assumption that maintaining yield with a decreased N supply rate can be beneficial for the environment and the fruit quality of tomato crop. Most of the knowledge on the N use efficiency (NUE), a genetic complex trait, comes from *Arabidopsis* and cereals. It is defined as the total biomass (or yield) per unit N supplied (Moll et al. 1982). NUE is divided in two main components: the nitrogen uptake efficiency (NUpE), defined as the plant ability to take up N from the soil, and the nitrogen utilization efficiency (NUtE), the plant ability to utilize (assimilate and transfer) N to the seeds (Xu et al. 2012). These two components involve several and interacting physiological traits including N absorption, translocation, assimilation, amino acid synthesis and catabolism, protein synthesis, sensing and signalling processes (The et al. 2021). In addition, some morphological traits are also playing a leading role in NUE. The best example is the introduction of the dwarfing genes in rice and wheat in the 1970s, which is arguably the largest improvement of the NUE achieved in crops (Liu et al. 2022).

Although considerable progress has been made for NUE understanding, relatively few studies have been performed in vegetables. In tomato, several genes involved in N-response have been identified through functional genomic studies. Reverse genetic approaches have been used to characterise several genes involved in N uptake (Fu et al., 2015), assimilation (Vallarino et al. 2022) or remobilization (Cao et al., 2022). In addition, many transcriptomic (Renau-Morata et al. 2021, Sunseri et al. 2023), proteomic (Xun et al., 2020) and metabolomic (Urbanczyk-Wochniak and Fernie, 2005; Larbat et al., 2014; Sung et al., 2015) approaches have identified differentially expressed genes and metabolic pathways involved in nitrogen metabolisms. Previous studies made progress in identifying nitrogen uptake efficiency (NUpE) QTLs in tomato. Several studies have characterised genotypes contrasting for NUE (Abenavoli et al. 2016, Rosa-Martínez et al. 2021, Aci et al. 2021) and proposed multiple physiological and molecular traits explaining these differences (*e*.*g*., root length and thickness, NO_3_^-^ influx rate, NO_3_^-^ storage, nitrate reductase activity, root cell electrical potentials, expression of NO_3_^-^ transporters). Rosa-Martínez et al. (2021) further characterised the variability and organoleptic qualities of a collection of ‘de penjar’ tomato varieties, identifying potential sources of resilience to low N fertilisation levels. However, none of these studies identified QTLs for N response. Two studies explored NUpE through linkage studies. Asins et al. (2017) used a population of RILs derived from *Solanum pimpinellifolium* as rootstock. They identified 62 significant QTLs and their results suggested a link between the content of hormones cytokinin and salicylic acid in roots and NUpE. Renau-Morata et al. (2024) studied a collection of 29 introgression lines resulting from a cross between the To-937 accession of *S. pimpinellifolium* and the *S. lycopersicum* cv Moneymaker and identified four candidate genes in the specific introgressed region associated with a greater photosynthetic capacity and biomass production under N deficiency conditions. Despite these studies, the overall identification of QTLs for N response in tomato remains notably limited.

The aim of this article is to investigate the genetic diversity and the genetic architecture of the response to long-term NO_3_^-^ deficiency in tomato. To this end, we characterised two large populations, an eight-way multi-parental genetic intercross (MAGIC) population and a diversity panel (core-collection of cherry type tomato accessions), to identify traits and QTLs linked to nitrogen response. These two populations offered the advantage of capturing a large genetic diversity, thus increasing the likelihood of identifying new genomic regions and candidate genes. Furthermore, a transcriptome analysis of differentially expressed genes in both fruit and leaves between NUE-contrasting genotypes was performed and allowed us to precise candidate genes under the QTLs. Overall, our findings offer new perspectives for understanding tomato plant responses to nitrogen stress condition and underline application in breeding programs.

## Results

### Impact of nitrate reduction on tomato

To assess the impact of limited NO_3_^-^ supply, two different panels of genotypes were studied: (i) a diversity panel (core collection, CC) of 143 small fruit accessions analysed by GWAS and (ii) an eight-way multi-parental genetic intercross (MAGIC) population composed of 228 lines derived from the cross of four large- and four small-fruited accessions, analysed using QTL mapping. Both populations were grown under two nitrogen conditions (2 and 10 mM NO_3_^-^, which we will refer to as the stress and control conditions, respectively), in the same greenhouse, during autumn and the following spring for CC and MAGIC, respectively. Several traits, including quality traits (fruit weight, soluble solid content (SSC) and pH), plant growth related traits (stem diameter, leaf length, sympode weight) and N-related traits (leaf Nitrogen Balance Index (NB, defined as the ratio of chlorophyll to epidermal flavonoids), nitrate petiole content, leaf nitrogen and carbon content) were measured. To explore how these traits responded to N reduction, phenotypic plasticity was quantified as the ratio between the mean values in both conditions at the genotype level. All the traits showed a large phenotypic variability in both panels (**Figure 1 - Supplementary figure S1**).

**Figure 1.**
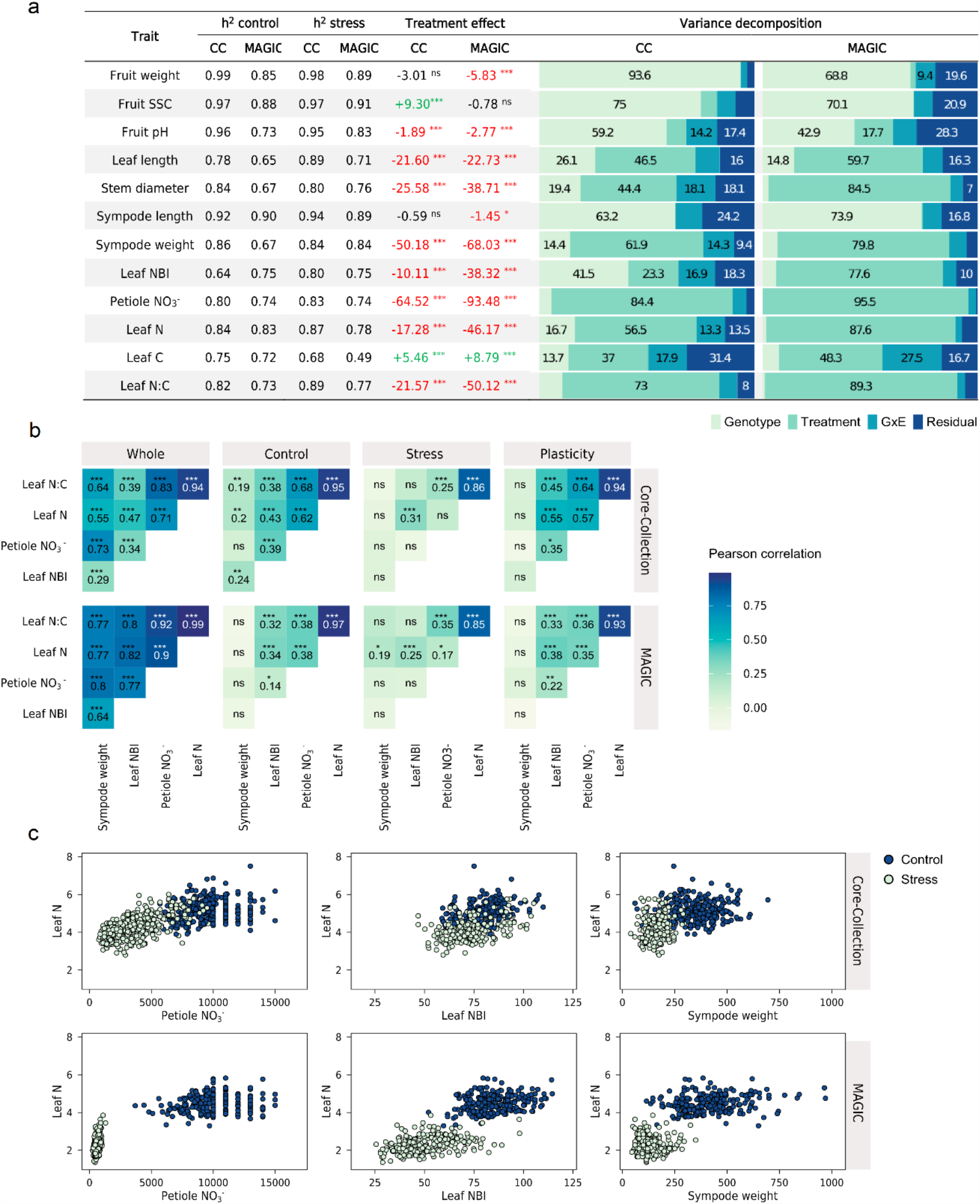
Impact of nitrate reduction on plant growth and N-related traits. Treatment effect (expressed as percentage of the difference between means in Stress and Control relative to Control condition) and variance decomposition (%) of 12 traits in the two genetic panels (CC, Core Collection; MAGIC, multi-parental genetic intercross population) grown in greenhouse under two N levels. SSC, *Soluble Solid Content*; Leaf NBI, *Leaf Nitrogen Balance Index*; Leaf N:C, *Leaf Nitrogen/Carbon content*. Significance of the *P*-value for the treatment effect: *** P < 0.001; ** 0.001 < P < 0.01; *0.01 < P < 0.05; NS, not significant. (**B**) Correlations between selected nitrogen-response traits at the whole level under low NO_3_^-^ (stress), high NO_3_^-^ (control) and plasticity conditions. (**C**) Details on the relationships between of Leaf N, Petiole NO_3_^-^, Leaf NBI, and Sympode weight.

Significant differences between treatments for each trait, except plant height, were detected. Other plant growth traits were largely negatively impacted by low NO_3_^-^ . Among these variables and for each panel, the plant weight was the most affected trait (on average -50.18% and -68.07% for CC and MAGIC panel, respectively). Fruit quality related traits (fruit weight, SSC and pH) were not strongly impacted by low NO_3_^-^ : the reduction in fruit weight was significantly different only in the MAGIC panel (CC: -2.75%; MAGIC -5.83%). Likewise, the increase in SSC was significantly different only in the CC panel (CC: +9.27%; MAGIC -0.78%). All the nitrogen related traits (NBI, petiole NO_3_^-^, leaf N, leaf C, leaf N:C ratio) were significantly impacted by the low NO_3_^-^, with nitrate content the most impacted variable (CC: -64.52%; MAGIC: -93.49%), as expected. Analysis of variance was conducted per each panel and trait (**Figure 1a**). Genetic variance was consistently higher in the CC panel. Furthermore, the treatment was higher than the genotypic effect for all the biomass and N-related traits. The broad sense heritabilities (h^2^) of the traits were moderate to high, ranging from 0.49 (Carbon content under low N) to 0.89 (Leaf N:C ratio for the CC under low N). Also, the *h*^*2*^ were consistent between conditions in most of the cases. Pairwise Pearson correlations (*r*) were calculated between pairs of traits within a treatment and for each trait between treatments (**Figure 1b**). In this last comparison, the correlations between traits were all significant (*p* < 0.05). However, the correlations were weaker between traits within each N condition. At low N, the correlations were higher compared to high N (presumably due to a narrower variability) in both panels. The highest correlation resulted between the variables petiole NO_3_^-^ and leaf N. The distribution of these two variables suggested a linear relationship under low NO_3_^-^ reaching a plateau under high N condition (**Figure 1c**). Finally, most of the significant correlations for plasticity traits were higher compared to low and high NO_3_^-^ conditions.

### Genetic dissection of N-related traits

Linkage analysis in the MAGIC population revealed 15 QTLs for N related traits. Among these, five were identified at high N (C), three QTLs at low N (S), and seven for plasticity traits (P), which represent the percentage of difference between the two conditions relative to the control condition. Notably, no significant QTLs were found for N content. Considering the overlapping confidence intervals as a single association, the analysis resulted in seven distinct QTL regions (**Table 1** and **supplementary figure S2**). The most significant QTLs for NBI under low and high N, as well as plasticity conditions (NBI:NO3_2) were observed on chromosome 2. Likewise, a QTL hotspot on chromosome 3 for petiole NO_3_^-^ content was detected across all the conditions (NO3_3). The confidence intervals (CIs) for QTLs ranged from 1.81 Mbp (NBI_4) to 59.1 Mbp (NBI_12). The overlapping confidence intervals and the similarity in founder haplotype assignment of phenotypic effects supported the hypothesis that these QTL clusters represented the same QTL. To reduce the candidate gene list, we applied a filter based on contrasts for the most different QTL founder allelic effects. This strategy significantly reduced the gene lists, with almost half reduction for NBI_4. However, for NBI_12 and NBI:NO3_2, more than 500 candidate genes remained. QTL analyses were also carried out on the agronomic and fruit quality characteristics (**Supplemental table S4**). As most of these traits (except plant weight) were not strongly influenced by nitrogen reduction, a large proportion of the QTLs detected were identified for both N conditions. In detail, 11 and 12 associations were detected under high and low N, respectively; 10 QTLs were shared between N conditions.

**Table 1.** QTLs detected under control (C), stress (S) and plasticity (P) conditions in the MAGIC panel for N related traits.

| Chr. | Position |  | LOD | Trait<sup>a</sup> | Cluster ID | Nb. genes<sup>b</sup> | Founders estimated allelic effects |  |  |  |  |  |  |  |
| --- | --- | --- | --- | --- | --- | --- | --- | --- | --- | --- | --- | --- | --- | --- |
|  | Lower | Upper |  |  |  |  | Ce. | Le. | Cr. | St. | Pl. | 1420 | Fe. | 0147 |
| 1 | 90,901,338 | 93,475,357 | 5.38 | Petiole NO<sub>3</sub><sup>-</sup> (P) | NO<sub>3</sub>_1 | 317 (260) | -0.53 | -0.71 | 0.29 | -0.40 | 0.80 | -1.60 | 0.59 | 1.56 |
| 2 | 33,973,517 | 36,931,627 | 5.33 | Leaf NBI (C) | NBI_2 | 240 (202) | -0.21 | 1.08 | -0.70 | -0.85 | -0.32 | -1.26 | 1.55 | 0.70 |
|  | 33,973,517 | 36,931,627 | 5.68 | Leaf NBI (P) |  | 240 (218) | -0.09 | 1.46 | 0.18 | -0.95 | -0.62 | -1.53 | 0.83 | 0.71 |
| 2 | 45,447,408 | 52,920,159 | 8.07 | Leaf NBI (S) | NBI: NO<sub>3</sub>_2 | 1 007 (708) | 0.37 | -0.52 | 1.57 | 0.37 | -1.19 | 0.62 | -1.46 | 0.25 |
|  | 45,447,408 | 55,336,340 | 9.54 | Leaf NBI (P) |  | 1 315 (1039) | -0.12 | 0.03 | 1.73 | 0.44 | -1.35 | 0.47 | -1.32 | 0.12 |
|  | 49,740,098 | 55,336,340 | 5.84 | Leaf NBI (C) |  | 743 (591) | -1.32 | 1.09 | 0.94 | 0.63 | -1.34 | 0.08 | -0.75 | 0.66 |
|  | 49,740,098 | 55,336,340 | 6.90 | Petiole NO<sub>3</sub><sup>-</sup> (C) |  | 743 (591) | 0.18 | 0.62 | 1.72 | -1.46 | -0.57 | 0.18 | -1.00 | 0.34 |
| 3 | 65,020,764 | 70,779,030 | 5.62 | Petiole NO<sub>3</sub><sup>-</sup> (S) | NO<sub>3</sub>_3 | 781 (261) | -0.52 | -0.34 | 0.32 | -0.01 | -1.52 | 0.09 | 2.04 | -0.05 |
|  | 65,831,575 | 70,779,030 | 5.47 | Petiole NO<sub>3</sub><sup>-</sup> (P) |  | 670 (158) | -0.92 | 0.13 | 0.33 | 0.82 | -1.95 | 0.25 | 1.21 | 0.14 |
|  | 68,052,409 | 70,779,030 | 5.39 | Petiole NO<sub>3</sub><sup>-</sup> (C) |  | 363 (146) | -1.00 | 0.85 | 1.04 | 1.00 | -1.45 | 0.63 | -0.78 | -0.30 |
| 4 | 60,952,746 | 66,467,941 | 5.97 | Leaf NBI (P) | NBI_4 | 693 (390) | -1.00 | 0.12 | 0.60 | 0.32 | -1.74 | -0.13 | 0.30 | 1.54 |
| 7 | 2,671,204 | 4,481,678 | 5.88 | Leaf NBI (C) | NBI_7 | 144 (129) | 0.15 | 1.11 | -1.79 | 1.38 | -0.42 | -0.12 | -0.59 | 0.28 |
| 12 | 4,892,313 | 64,046,622 | 8.38 | Leaf NBI (S) | NBI_12 | 1 414 (1145) | 1.25 | -0.13 | -0.30 | -0.69 | 1.83 | -0.88 | -0.27 | -0.81 |
|  | 5,047,381 | 64,046,622 | 7.83 | Leaf NBI (P) |  | 1 401 (1135) | 1.56 | -0.06 | -0.45 | -0.88 | 1.53 | -0.75 | -0.13 | -0.82 |
| 12 | 9,476,793 | 64,125,346 | 5.21 | Petiole NO<sub>3</sub><sup>-</sup> (P) | NO<sub>3</sub>_12 | 1 222 (974) | 1.80 | -0.56 | -1.06 | 0.70 | -0.08 | -0.35 | 0.66 | -1.12 |

**a:** Trait nomenclature: Trait\_Condition; **C:** Control; **S:** Stress; **P:** Plasticity.
**b:** In parentheses the number of genes selected by allelic contrast.
**Ce:** Cervil, **Le:** Levovil, **Cr:** Criollo, **St:** Stupicke Polni Rane, **Pl:** Plovdiv, **1420:** LA1420, **Fe:** Ferum, **0147:** LA0147

GWAS analysis in the CC panel revealed 12 QTLs for N-related traits (**Table 2** and **supplementary figure S3**). Among these, one and seven associations were identified for high (C) and low N (S) conditions, respectively; while 12 associations were related to plasticity traits (P). By contrast to the MAGIC linkage analysis, the GWAS approach yielded only three colocalized QTL clusters on chromosomes 6, 11 and 12. To determine the number of candidate genes, we filtered for 10 kb window around each significant marker (not only lead SNP). Consequently, the number of candidate genes per QTL was significantly reduced compared to the MAGIC panel, ranging from 0 to 9 genes. GWAS analyses for fruit quality and agronomic traits were also conducted (**Supplemental table S5**). As observed in the MAGIC, a large number of QTLs were identified in both conditions: 18 and 16 associations were found under high (C) and low N (S), respectively; 15 QTLs were shared between N conditions.

**Table 2.**
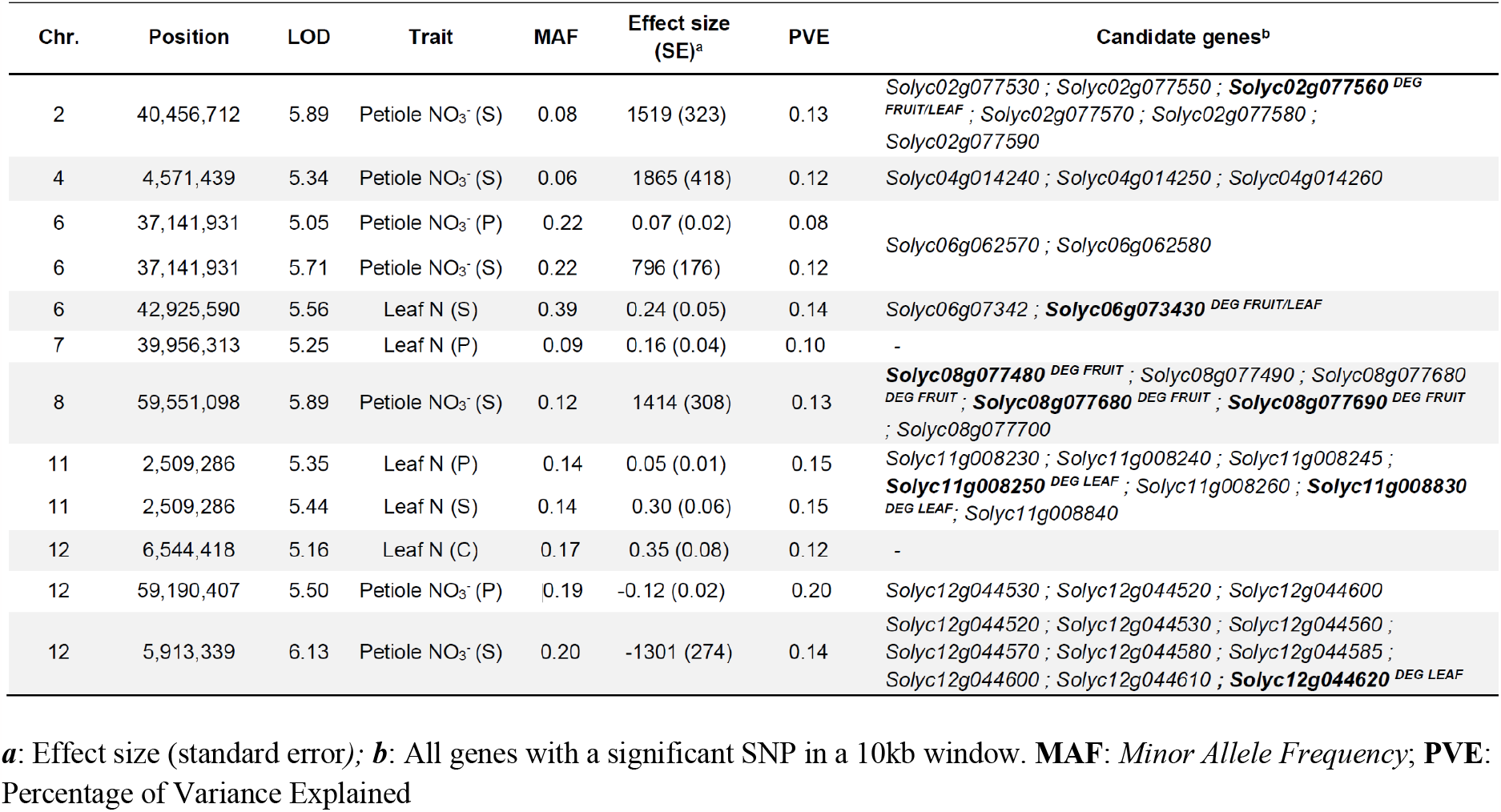
QTLs detected for petiole NO_3_^-^ and leaf N content by GWAS in the CC panel.

### Transcriptomic analysis of N-responsive genes

The transcriptome profiles of four tomato accessions were analysed under the same contrasted N-regime (2 and 10 mM NO_3_^-^). The accessions Ferum, Levovil, Cervil and LA1420, showing the most contrast for fruit weight and plant vigour (thus expected to differ for N response), were chosen among the parents of the MAGIC population. The transcriptomes were analysed on leaves and fruits. The RNA sequencing firstly yielded 26,544 and 25,414 expressed genes in leaves and fruits, respectively, resulting in 15,264 and 12,827 genes after filtering for sufficient and consistent expression across the samples. Principal Component Analysis (PCA) for differential gene expression was conducted on normalised counts (**supplementary figure S4 & S5**). The first two components of the PCA accounted for 41,4 % and 40,6 % in fruit and leaves, respectively. The variation was mainly related to genotypes and/or treatments while no effect of the replicates appeared. The differences between genotypes were clearer on fruit while the treatment impact was more significant in the leaves. The higher number of DEGs was identified between genotypes regardless N-treatment. Indeed, in leaves, this number ranged from 4,465 (Ferum control *vs* Levovil control) to 8,906 (Ferum *vs* LA1420) with an average of 6.818 DEGs (**supplementary figure S6**). At high N (Control, C) the small-fruited accessions showed contrasting expression profiles, at low N (S) they tended to exhibit similar expression responses, as opposed to the big-fruited accessions. In fruit, the number of DEGs ranged from 5,171 (Ferum stress *vs* Levovil stress) to 9,740 (Cervil *vs* Levovil) with an average of 7,088 DEGs. At high N (7,741 DEGs) and low N (8,543 DEGs) the most contrasted genotypes were Cervil and Levovil. The least contrasted genotypes were LA1420 and Levovil at high N (5,236 DEGs), Ferum and LA1420 at low N (5,171 DEGs). The number of DEGs between N-treatments were lower than the comparisons between genotypes. About four thousand (4,056) and 6,526 DEGs were found in response to N treatment in fruit and leaves, respectively, among which 1,677 (19%) were shared between tissues. In the comparison between N treatments (C vs S), 4,752 and 2,405 DEGs were detected in leaves and fruit, respectively when considering all the genotypes. Among these genes, 3,628 (76.3%) in leaves and 1,717 (71.4%) in fruit were also found at least once in a genotype-specific contrast (*e. g*. C vs S in Levovil only). Then, they could be considered as genotype specific, the significance of the differential expression being supported mostly by the results of a single genotype (**Figure 2a & 2b**). Among the 4,752 DEGs found in leaves, 1,124 were detected in the C *vs* S comparison, regardless of the genotypes. The remaining DEGs were genotype-specific in the same comparison. The higher number of DEGs were found in big-fruited accessions (4,213 DEGs), while only 313 DEGs were specific to the small-fruited accessions. Finally, 71 genes were found differentially expressed in all the five comparisons. It was observed in the PCA that leaves were more sensitive to the low N and fruit to genotype differences. The same trend was observed when considering the number of DEGs by tissue and by contrast. When focusing solely on DEGs identified in the C *vs* S comparison but gathered by accession, more DEGs were detected in big fruited tomatoes, 3,982 DEGs in Ferum leaves and 2,372 DEGs in Levovil leaves, while in the small-fruited accessions, 559 DEGs were detected in Cervil leaves and 446 in LA1420 leaves. The difference between accessions was reduced in the fruit but followed the same pattern. Roughly the same number of genes were differentially expressed in the small-fruited accessions but the number of DEGs decreased in big-fruited accessions (with 1,460 DEGs in Ferum and 1,555 DEGs in Levovil).

**Figure 2.**
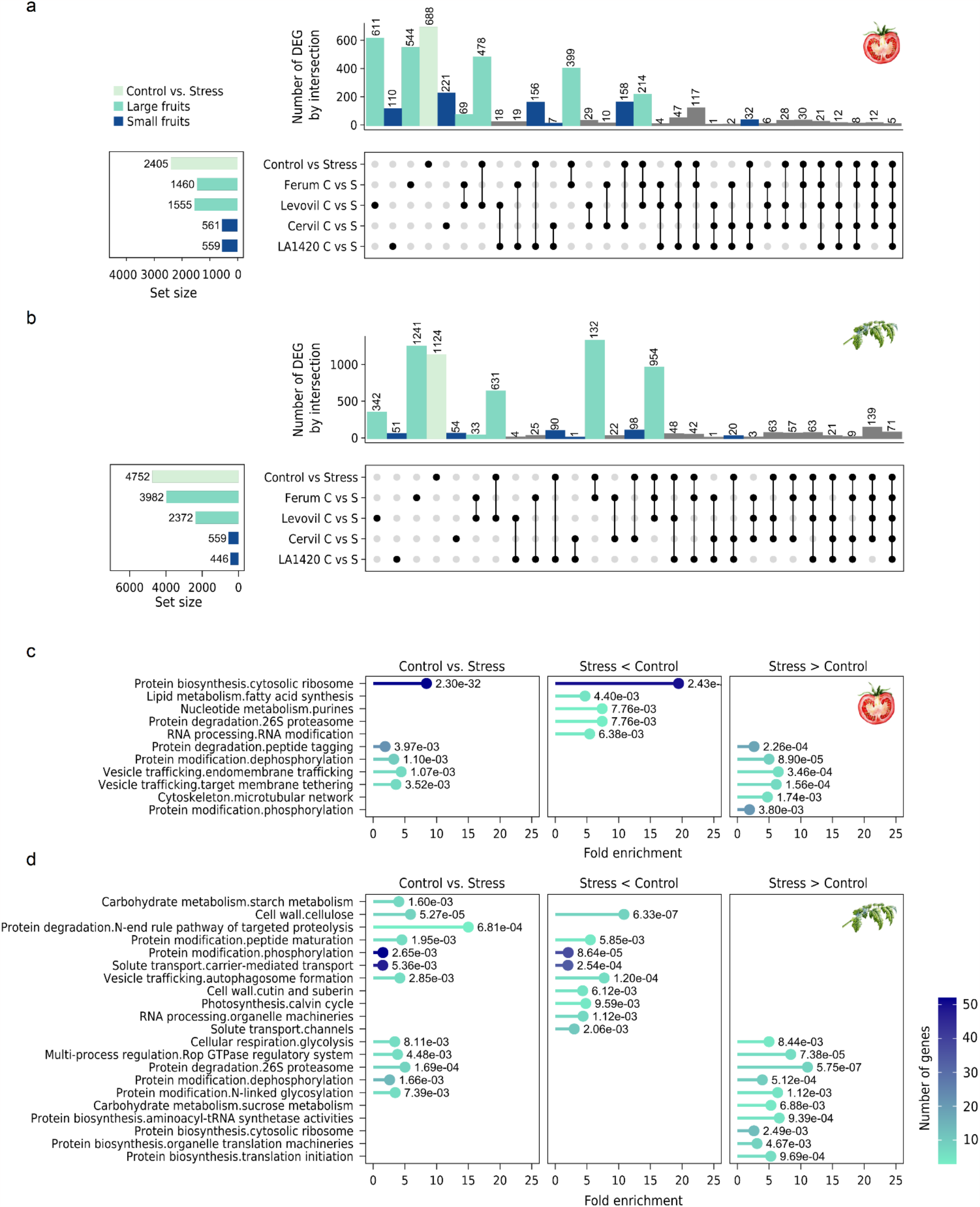
Differential Gene Expression and Enrichment Analysis in leaves and fruits. Panels A and B are upset plots detailing the number of DEGs for each contrast, as represented by the left side barplot, in fruit (A) and leaf (B). The barplot illustrates the count of shared DEGs across contrasts, with combinations depicted in the central grid. The ‘Control vs Stress’ contrast encompasses all accessions, while other contrasts highlight treatment responses specific to individual accessions. Panels C (fruit) and D (leaf) present enrichment results based on the second pathway levels as defined by the MapMan classification. The y-axis lists pathways with at least one identified DEG, while the x-axis represents the log-fold change between the two conditions under consideration. Point colour corresponds to the number of genes identified within a pathway. P-value of the enrichment test is also displayed.

To obtain an overview of the biological pathways regulated in response to N-treatments, a gene enrichment analysis based on the MAPMAN classification was first performed on the 1,124 DEGs (**Figure 2d**) detected in leaves for all genotypes. Twelve pathways were found enriched. Among these pathways, most genes belonged to the ones corresponding to protein modification and degradation. Vesicle trafficking and solute transport were also enriched as well as cellular respiration, carbohydrate metabolism and cell wall. Focusing on the 511 DEGs up-regulated in N-stress condition, protein modification, degradation and biosynthesis pathways remained enriched as well as cellular respiration and carbohydrate metabolism. For the 613 DEGs down-regulated in N-stress condition, the protein modification and degradation pathways were still enriched together with the vesicle trafficking and solute transport pathways. Interestingly, for the “protein modification” pathway, the genes of the sub-pathway “protein modification.phosphorylation” were down-regulated in N-stress condition while the genes of the sub-pathway “protein modification.dephosphorylation” were up-regulated. The same analysis was performed on DEGs detected in fruit (**Figure 2c**). Although only 688 DEGs were considered, five pathways were enriched. At the first level of classification, the same pathways were found enriched (“Protein biosynthesis”, “Protein degradation”, “Protein modification”, “Vesicle trafficking”). The only pathways shared between tissues were “Protein biosynthesis.cytosolic ribosome” and “Protein modification;dephosphorylation”. In fruit, “Protein biosynthesis.cytosolic ribosome” was also enriched for genes down-regulated in N-stress condition, while in leaves it was enriched for down-regulated genes. Finally, a focus on genes involved in nitrogen metabolism through a list of 185 genes established based on published data and annotation (**supplementary table S7**) was performed. Among these 185 genes, 33 and 53 were found differentially expressed in the present experiment, in the fruit and leaves, respectively. Their average expression profiles across the eight samples are presented in **Figure 3**. The clustering of samples confirmed that the differences was more important between genotypes rather than N treatments in fruit. The clustering of DEGs matched roughly with gene families, a first cluster corresponding to Amino acids biosynthesis, a second one to Autophagy / senescence and Nitrate reductases and the last cluster to Amino acids transporters. Most of the genes presented a positive logFC or if negative close to zero, with the exception of Solyc01g080280 (logFC = -3.43) in Ferum and three genes in Levovil: Solyc04g077050 (logFC = -2.56), Solyc07g051950 (logFC = -2.27) and Solyc06g060110 (logFC = -2). These genes were mainly expressed in N stress condition. In leaves, the candidate genes showed a specific profile in Ferum under N-stress, by contrast to LA1420 under control condition and fairly different compared to Ferum under control. Three main gene clusters appeared differentially expressed, in the first, up-regulated genes related to autophagy and senescence under control were included. In the second cluster, genes up-regulated under control condition related to the Amino acid transporters. The last cluster gathered genes from the same class but less expressed in stress conditions.

**Figure 3.**
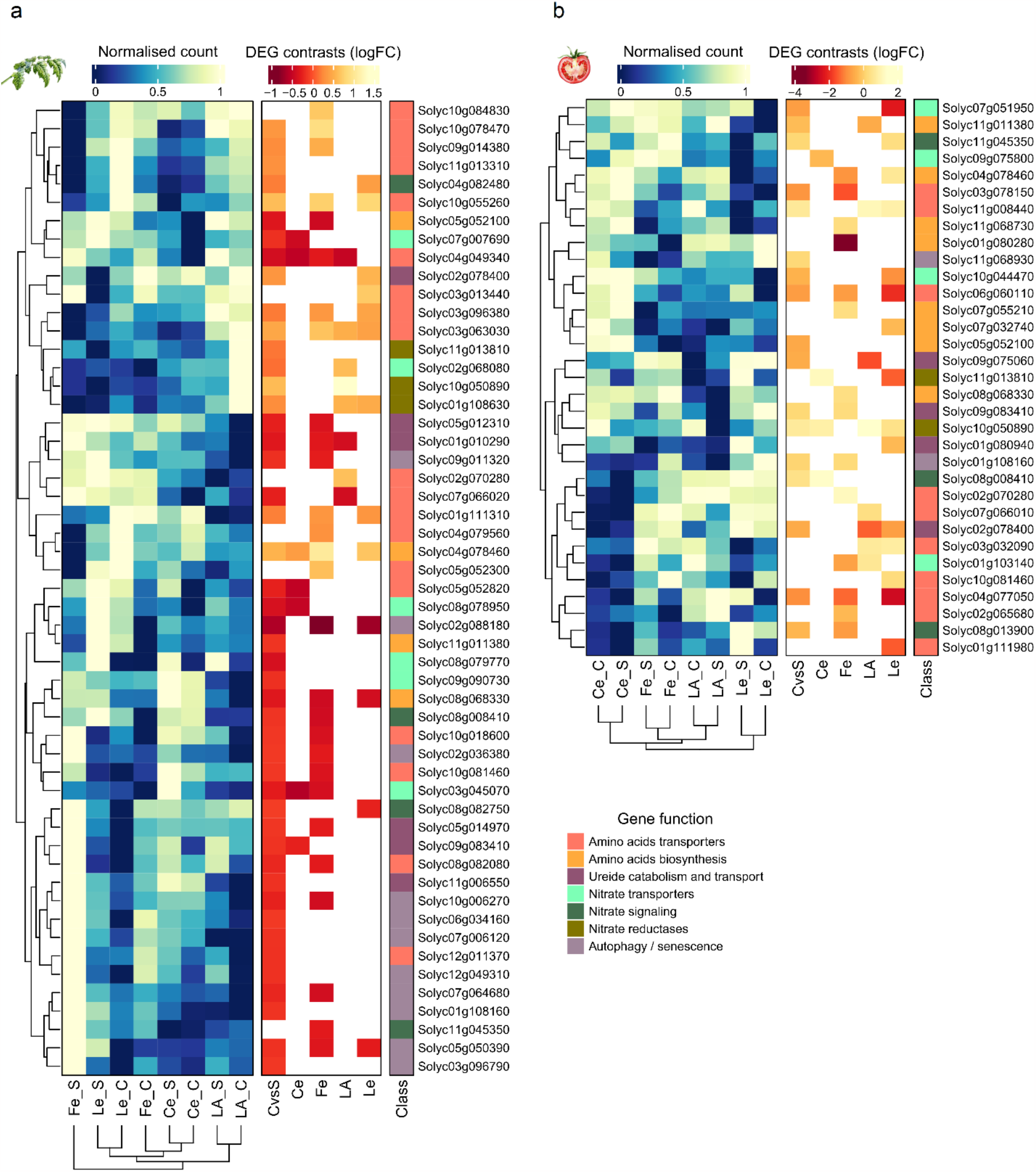
Expression profile and results of differential expression for genes involved in N metabolism and response to N stress in leaf (a) and fruit (b) Heatmap of the normalised count averaged over the samples for each combination of accession x treatment, for each of the genes found expressed either in the fruit (right panel) or the leaf (left panel). The genes are gathered and coloured based on their class and function. The right panel shows the contrasts in which the genes were found to be differentially expressed, the colour level corresponds to the log fold change between the tested conditions. A negative log fold change means that the gene is more expressed in stress conditions. Fe: Ferum, Le: Levovil, Ce: Cervil, LA: LA1420, C: Control condition, S: Stress condition.

### Candidate gene identification

In order to identify candidate genes for the QTLs, we first focused on QTLs that included less than 500 candidate genes in their confidence intervals for the analysis of genes derived from MAGIC. Consequently, two clusters (NBI:NO3_2 and NBI_NO3_12) were excluded from subsequent analyses. To identify potential candidate genes, we focused on candidate genes included in the QTL intervals and differentially expressed between N treatments (C *vs* S). Following this strategy, we significantly narrowed down the list of candidate genes (**Figure 4**; list available in **Supplementary Table S6**). Most of the identified candidate genes were associated with nitrogen remobilization functions. Among them, two genes were involved in senescence and autophagy (*Solyc01g104080, Solyc04g076720*), as well as three nitrogen compound transporters (*Solyc02g065680, Solyc07g008440, Solyc07g008520*). The list of candidate genes identified by GWAS is more limited compared to linkage mapping QTLs. Indeed, only 35 genes were identified closed (within a 10kb window) to significant SNPs for N related traits (**Table 2 and supplementary Table S7**). Among these, five genes showed differential expression in leaves, five in fruit, and two genes were differentially expressed in both tissues. Notably, within the significant DEGs identified at the chr02_NO3_S QTL, *Solyc02g077560*, that codes for the Auxin Response Factor 3 (*SlARF3*), emerged as the most interesting candidate gene. More interestingly, it was the only gene among the candidates associated with SNPs in the chr02_NO3_S QTL that exhibited differential expression in both tissues. The comparative sequence analysis among genotypes was not able to identify any polymorphism in the coding region. Thus, the variation in gene expression was probably due to a polymorphism in the regulation sequences. In *Sl*ARF3 RNAi mutants, substantial morphological changes were observed, including a decrease in the density of epidermal cells and trichomes in leaves, as well as inhibited xylem tissue development (Yifhar et al. 2012). Furthermore, cis-eQTL analysis revealed the presence of regulatory variants influencing its expression at the fruit level (Zhu et al. 2018).

**Figure 4.**
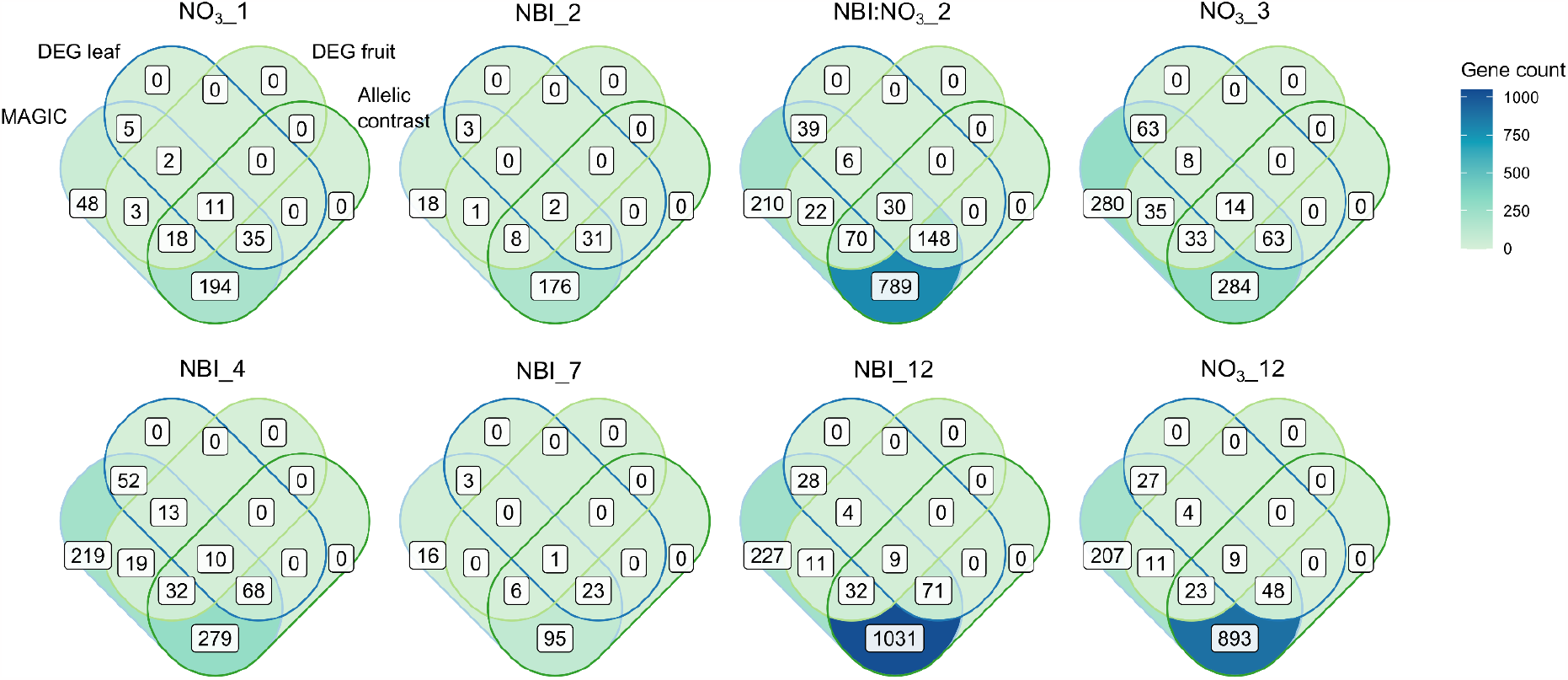
Intersection of numbers of candidate genes from linkage mapping in the MAGIC population and differentially expressed genes (DEG) between stress and control nitrogen treatments in leaf (DEG leaf) and fruit (DEG fruit) MAGIC: number of genes found in the QTL confidence interval; Allelic contrast: genes found within the confidence interval and whose polymorphisms correspond to the allelic effect estimated by the founder lines.

We identified several other promising candidate genes associated with different QTLs: *Solyc08g077480* encoding for a SnRK1-interacting factor and belonging to the senescence-associated family protein. It was also found to be differentially expressed in fruit; *Solyc11g008830*, an homolog of *A. thaliana* ASYMMETRIC LEAVES2 whose mutant is involved in the generation of leaf lobes and leaflet-like structures (Capua and Eshed 2017); *Solyc12g044610*: also known as MYB1R1 or MYBS3, encodes a stress-responsive MYB transcription factor. It is upregulated in response to cold conditions (Guo et al. 2022) and low phosphate levels.

## Discussion

### Phenotyping NUE-related traits in tomato

The assessment of NUE *stricto sensu*, i.e., net dry mass production per unit of N uptake, in indeterminate (vine) tomato plants is more complex than in other crops such as cereals as it is a long-cycle crop harvested multiple times. Thus, this crop is subject to an export of biomass over time, in both fruit and leaf biomass (towards senescent leaves and lateral shoots which are removed). We thus chose to report an index of biomass production per unit of internode as N-response biomass trait. Furthermore, single trait cannot unravel the intricacies of NUE genetic control, regulated by several pathways and physiological mechanisms. Therefore, we targeted other N related traits including leaf N and C content, petiole NO_3_^-^ and used Dualex Leaf-Clip to assess the NBI. Correlation analysis showed that there was no strong correlation between leaf nitrogen content and other nitrogen-related traits, and therefore dismissed the idea of strong pleiotropy between these traits. The correlation between leaf N content and NBI was weak to moderate for both nitrogen treatments and panels (*r* = 0.25-0.43). The large diversity of leaf structure within the two panels of genotypes could probably explain this moderate relationship. Besides, several studies have already reported the relationship between chlorophyll-based measurements (NBI) and leaf N content is cultivar dependant (Minotti et al. 1994, Monostori et al. 2016, de Souza et al. 2020). The petiole NO_3_^-^ content was only significantly correlated with the leaf N content under low N condition. This relationship could be explained as a luxury uptake of N. At high N condition, significant correlations between N-traits were not observed because they were at the plateau of the curve (**Figure 1c**). Surprisingly, the correlation between sympode weight and N content was not significant (except at low N condition in the CC). Renau-Morata et al. (2024) reported the same result using a *S. pimpinellifolium* introgression line panel. In addition, as mentioned above, this measure is probably too biased by biomass exports during tomato life cycle and the difference in senescence rate observed between genotypes. To get a true estimate of biomass, it would be more accurate to switch to non-destructive imaging techniques. The analysis of NUE could then be similar to that described for potato, by analysing canopy development parameters (Khan et al. 2010). This type of approach is particularly suitable for drone phenotyping and seems to work well even for wild genotypes with indeterminate development in the field (Johansen et al. 2020). Finally, the characterization of senescence rate in relation to nitrogen remobilization is a promising trait to be investigated.

### Transcriptome insight into N deficiency

The transcriptome analysis of four tomato accessions grown under contrasting N supply did not highlight a specific biological pathway strongly regulated, whether the analysis was performed without *a priori* through enrichment or by focusing on candidate genes. Nevertheless, the biological pathways found enriched when considering DEGs identified in response to the N-treatment were involved in protein biosynthesis, modification and degradation. These pathways were found enriched both for genes up- and down-regulated under N stress condition. The vesicle trafficking and solute transport pathways were also found significantly enriched mainly in the down-regulated DEGs under N stress condition. The different N rates supplied induced variability in the transcriptome across the four genotypes under study. However, this variability was lower under N-stress compared to the variation identified between genotypes regardless N treatment. The contrasting pattern of gene expression in the fruit between Levovil and Cervil were expected due to their smallest and largest fruit weight and plant vigour. The other similarities and differences in gene expression were not easily explained by morphological traits. Indeed, Ferum and Levovil were expected the most similar accessions for their responses in all the conditions, because they were both big-fruited tomatoes and similar for many morphological traits, but also highly convergent when comparing their sequence polymorphisms against the reference genome (Causse et al. 2013). The specificity of genotype response and the prevalence of genotype effect over treatments emphasises the necessity to follow several accessions in physiological and genetic analysis to obtain the most complete possible overview of N use efficiency for a species. A buffering effect between leaves and fruits was observed for the number of DEGs identified in response to N-treatment. It was not the case for DEGs identified between genotypes regardless N-treatment. Such buffering effect in fruit was already described for small-fruited accessions in response to water deficit(Albert et al. 2018), but this effect seemed stronger in our experiment, with a similar number of DEGs between leaf and fruit for the small-fruited accessions and a significant decrease for the big-fruited accessions. Nevertheless, despite this low number of DEGs in fruit, it was still expected to induce phenotypic differences at the fruit level in response to N stress. By contrast, the impact of N stress on fruit, albeit significant, was limited compared to the effect on plant biomass production. This indicates that in stress conditions, fruit growth was maintained at the expense of vegetative growth which induced specific regulation.

The current knowledge of plant response to low N conditions at the expression level have been mostly obtained in the model plant *Arabidopsis thaliana* (Krapp et al. 2011, Balazadeh et al. 2014, Luo et al. 2020) or row crops such as durum wheat (Curci et al. 2017), wheat (Zhang et al. 2021), rapeseed (Ahmad et al. 2022), or rice (Shin et al. 2018). However, limited information is available on vegetables such as tomato. It is even more scarce when looking for RNAseq analysis. Two recent studies tried to tackle this question. The transcriptome response of 35-days plant of the cultivar Moneymaker was assessed when cultivated with a solution containing 4 mM (stress) or 8 mM (control) of nitrogen (Renau-Morata et al., 2021). The other study focused on the short term (8h - 24h) transcriptome response differences in shoot and root between a pair of high-NUE (Regina Ostuni) and low-NUE (UC82) cultivars (Sunseri et al., 2023). The plants were grown for 20 days and then stressed (0.5 mM N) or maintained in controlled conditions (10 mM). By comparison, the first study focused on only one genotype in less stressful conditions while the second study applied a stronger but shorter stress. Despite these differences between our protocol and that from Renau-Morata et al. (2021), shared DEGs between the experiments were identified (**supplementary figure S7**). Indeed, Renau-Morata and colleagues highlighted 257 and 1,440 DEGs in roots and leaves (61 were shared between tissues), respectively, 31.5 % (81) and 49.5 % (713) of which were respectively also detected in leaves in the present study. Overall, in four sets of DEGs, leaves and fruits in the present study, roots and leaves in Renau-Morata et al. (2021), 12 DEGs were shared among all the sets (**supplementary figure S7**). The higher overlap between studies was found in leaves with 471 shared DEGs. Our results appeared consistent due to the relatively high overlap between comparable organs, however the present study detected more DEGs. Furthermore, Renau-Morata and colleagues identified a specific role of the alternative respiration and chloroplastic cyclic electron transport in tomato, not observed in other crops. The alternative oxidase respiration (AOX) consumes sugars and starch in excess in leaves and is involved in the plant C/N balance under N stress (Noguchi 2006). In our experiment, the pathway “Carbohydrate metabolism.starch metabolism” was significantly enriched in the DEGs identified in leaves for the N-stress response. The pathways “Carbohydrate metabolism.sucrose metabolism” and “Cellular respiration.glycolysis” were significantly enriched in the DEGs up-regulated in N-stress condition, in all the genotypes. Only one of the genes specifically associated with the AOX pathway was sufficiently expressed to be analysed (*Solyc08g075540*) and was found down-regulated in the control (high N) in fruit (C *vs* S, Cervil C *vs* S and Levovil C *vs* S). The three genes related to the chloroplastic cyclic electron flow (CEF) pathway (*Solyc08g007770*; *Solyc09g090570*; *Solyc08g080050*) were differentially expressed. The first two genes were down-regulated in fruit, while *Solyc08g080050* was also down-regulated in leaves. However, at low N, *Solyc09g090570* was up-regulated in fruits and down-regulated in leaves.

Sunseri et al (2023) identified 297 and 154 DEGs in response to N treatment in shoot and root, respectively, which must be compared to the 923 (154) DEGs for the genotype by treatment interaction, 5,102 (3,800) DEGs for the time x treatment interaction and finally 5,480 (4,054) DEGs for the genotype x treatment x time interaction. Overall, they identified 7,667 and 6,015 unique DEGs in shoot and root, respectively. In this study, they identified a number of DEGs consistent with our study but their results also underlined the importance of considering the kinetic of gene expression (Sunseri et al. 2023). One of the hypotheses for our results might be that the long duration of the stress ended homogenising the plant’s responses or allowed compensation by other unknown mechanisms. Interestingly, both studies highlighted the role of nitrogen transporters. In our study, the high-sensitive N transporter (*NRT2*.*1, NRT2*.*2, NRT2*.*3* and *NRT2*.*4*) were not highlighted as DEGs, most probably because at 2 mM we were already outside of N range of NRT2 gene family activity. However, at low N, the low affinity transporter *NRT1* (*Solyc08g078950*) was up-regulated in leaves for all the genotypes. Other transporters were also found as DEGs in leaves: *AMT1*.*1* (*Solyc09g090730*), *AMT1*.*3* (*Solyc03g045070*), *NF-YA5* (*Solyc08g062210*) and *NF-YA9* (*Solyc01g008490*), all being down-regulated at low N.

Thus, several studies studying different genetic stocks and methodologies highlighted shared regulated genes which might consist in a common basis of regulation. However, the differences on plant genetic origin and in N stress application and duration induced variation in the direction of regulation (up- and down-). Furthermore, the majority of DEGs were specific to the study indicating that tomato plant response to limiting N conditions requires fine tuning and depends on a multiplicity of mechanisms.

### QTLs analysis: low pleiotropy for N-related traits and candidate genes

The genetic architectures of the nitrogen related traits exhibited a marked reliance to specific mapping population adopted, based on the different quantitative trait loci (QTLs) identified between the populations. This observation aligns with previous findings that reported similar outcomes for different traits within the same populations (Pascual et al. 2016, Bineau et al. 2021). The detection of population-specific QTLs probably arises from different reasons. Firstly, the two populations show significantly different allelic compositions as they were composed to represent different genetic groups. For instance, the CC panel encompasses accessions from diverse geographical origins (Albert et al., 2016) and spans three unique genetic groups (SP, SLC, and mixture) covering the domestication stage of tomato evolution. Conversely, the MAGIC population segregates for alleles coming from eight very diverse cultivated lines derived from later breeding efforts, with four cherry and four large-fruited accessions. Thus it represents the allelic diversity of improved tomato breeding materials, as proposed by Lin et al. (2014). Secondly, the two populations were grown at different seasons and the stress conditions may have correspond to slightly different quantities of N received per plant, although the concentrations were the same. Beyond the inter-population disparities, a limited number of QTL clusters influencing multiple traits were detected, especially within the MAGIC panel. Notably, we pinpointed QTLs exerting influence across the treatments (e.g., NBI:NO3_2 and NO3_3). Each trait might be modulated by a distinct ensemble of genes or genetic variants. Hence, the non-convergence of QTLs may be indicative of trait-specific genetic regulation rather than a lack of genetic influence.

Nevertheless, the MAGIC and the CC panels provided lists of candidate genes of drastically different sizes due to the different numbers of SNP markers between the two populations and LD structures. The intersection of the lists of candidate genes retained in the confidence intervals with that of DEGs derived from the RNAseq experiment should be interpreted with caution. Indeed, the choice of tissues used for RNAseq has a considerable impact on the results obtained. For instance, the expression of nitrate transporters (NRT) is negligible in fruit and leaf whereas these genes are particularly important for nitrate uptake. Thus the spotlighted candidate genes were mainly related to the N Utilization Efficiency (NUtE) component, such as nitrogen remobilization (*Solyc01g104080, Solyc04g076720, Solyc08g077480*), transport of nitrogen compounds (*Solyc02g065680, Solyc07g008440, Solyc07g008520*) or involved in morphogenesis (*Solyc02g077560, Solyc11g008830*). These genes emerged as potential candidates for functional validation.

## Conclusion

In this study, several nitrogen related traits were analysed in tomato under greenhouse experiments. Our results revealed a large range of responses to reduced N supply, albeit the genomic interactions with N treatment appeared limited. The QTLs identified for these traits exhibited little colocalization, suggesting that multiple genes underlie the genetic diversity of response to low N availability. In addition to QTL analysis, we conducted a differential gene expression experiment in fruit and leaves based on the contrasting N supply and revealed a large number of genes impacted by N level. The impact of genetic background on the gene expression was also underlined. The intersection analysis between genes included in the most significant QTLs and the identified DEGs allowed to select few genes differentially expressed located in the QTL regions. This analysis highlighted new candidate genes as key players in nitrogen usage efficiency, mainly in the NUtE component. The present study confirmed the complex genetic architecture that governs NUE -related traits in tomato. Our results offer a promising set of candidate genes for breeding NUE-enhanced tomato varieties. Overall, harnessing genomic regions associated with N-related traits in tomato will contribute to the establishment of appropriate breeding schemes for an early selection of low N input -adapted genotypes. The functional validation of candidate genes identified in this work will contribute to their characterization.

## Materials and methods

### Plant materials and growth conditions

Two panels were studied to analyse the genetic response to low N input: a core-collection (CC) of 143 cherry-type tomato genotypes and a collection of 228 lines from an eight-parent multi-parental genetic intercross (MAGIC) population. The CC panel is slightly modified compared to that described by Albert et al. (2016) (**Supplementary table S1**). It is composed of 115 genotypes from *S. lycopersicum* var. *cerasiforme* (SLC), 9 genotypes from *S. pimpinellifolium* (SP), and 19 admixed genotypes. The MAGIC panel is a subset of 228 lines of the population introduced and described by Pascual et al. (2015).

The CC and MAGIC populations were evaluated in the same greenhouse in Avignon INRAE Research Centre (Avignon, France) from August to November 2021 and March to July 2022, respectively. Each trial was conducted with the same culture conditions as described below. For each panel, 800 plants were grown, and two treatments were applied: optimal N (control) and N limited conditions (N stress). The experiments were carried out in rockwool-based hydroponic conditions. The nutritive solution included 10 mM and 2 mM NO_3_^-^ for the control and the N stress condition, respectively. It was provided by drip irrigation using a Nutricontrol Minimac A® (Nutricontrol, Spain) fertigation equipment. A constant nutrient concentration and conductivity (EC) were maintained in the hydroponic solution throughout the experiment. In the stress condition, Nitrate was substituted by Sulfate.

The experimental design included two and three replicates per genotype for the control and N-stress condition, respectively in the CC panel. In the MAGIC panel experiment, the eight parents of the population and their four F_1_ hybrids were used as controls and replicated twice. In addition, each of the 228 recombinant lines was replicated once (156 lines) or twice (72 lines) in the control treatment, and all the lines twice in the low N treatment.

### Phenotyping

Both panels were phenotyped for phenological, morphological, nitrogen-related and fruit quality traits. The morphological traits included stem diameter, leaf and sympode length and weight (measured between truss 3 and 6). To assess plant N status, different traits were recorded on the same samples: NO_3_^-^ petiole content, leaf nitrogen and carbon content. Petioles were frozen into Bioreba® extraction bags (Bioreba, Switzerland) and stored at −20°C until the analyses. Petiole sap was extracted by squeezing the sample using a paddle lab blender. NO_3_^-^ content was recorded using an ionometer (LAQUAtwin NO3-11C, Horiba Scientific, Japan). Before each measurement series and after every 15-25 samples, a two-point calibration was conducted using the 150 and 2000 mg NO_3_^-^.L^-1^ standards provided by the manufacturer. All the samples from the control condition were diluted 10 times with demineralized water to maintain concentrations under 1500 mg.L^-1^ to avoid underestimation of NO3--N content (Peña-Fleitas et al. 2021). Leaflets were dried in a forced-air oven (70°C) during 48:00 h before grinding. N and C contents of each sample were measured from 5 mg of powder using an auto-analyser (Flash EA 1112, Thermo Firmingam Milan, Italy). Three harvests (three to ten fruits each according to the fruit size) of red ripe fruits were conducted, from the 3^rd^ to the 6^th^ truss, for the analysis of fruit quality traits. The fruits were pooled by genotype and harvesting time to measure fruit weight. Then, crushed fruit pericarp was used to measure soluble solid content using an electronic refractometer (Atago PR-101, Atago Co. Ltd, Japan), and pH using a potentiometric titrator (TitraLab AT1000© series, HACH Company, USA). Phenotypic data are provided in **Supplementary Table S2 and S3**.

### SNP calling, raw data filtering

Seventy-two (72) out of 143 CC genotypes were already sequenced and the data are available at the NCBI platform (**Supplementary Table S1**). Sequence quality was examined with V0.11.8 (https://github.com/s-andrews/FastQC) and trimmed with fastp 0.20.0 (Chen et al. 2018) with the following parameters: max_len1 350, cut_mean_quality 20, cut_window_size 4, complexity_threshold 30. For each genotype, fastq files from several libraries were merged if available, and they were aligned on the reference genome Heinz_1706 v.4.0.0 using bwa 0.7.17, PCR duplicates were removed with SAMtools v1.9 (Li et al. 2009). Variant calling was performed by gatk4 v4.1.4.1 (https://github.com/broadinstitute/gatk).

The remaining CC genotypes were newly sequenced. Genomic DNA was extracted from 3-week-old plants using the DNeasy Plant Mini Kit (Qiagen, Hilden, Germany) following the manufacturer’s instruction. The amount of DNA was quantified using a Qubit fluorometer (Invitrogen, Carlsbad, CA, USA) and the 260/280 and 260/230 ratios were assayed using the NanoDrop 1000 spectrophotometer (ThermoFischer, Waltham, MA, USA). DNA samples were sent from ENEA (Casaccia Res. Ctr., Rome, Italy) to NOVOGENE for sequencing (10X depth). Finally, all the genotypes were combined in a single database and the SNP extracted (filter parameters QUAL < 30.0, QD < 2.0, FS > 60, MQ < 40, SOR > 3, MQRankSum < –12.5, ReadPosRankSum <–8 and depth <3.). The whole pipeline was implemented in Snakemake v5.8.1 (Köster and Rahmann 2012) and the containers were built for all the software using Singularity v3.5.3 (Kurtzer et al. 2017). SNPs with minor allele frequencies (MAF) <0.05 and call rates <80% were discarded using PLINK 2.0 (https://www.cog-genomics.org/plink/2.0/). After filtering, missing genotypes were imputed using BEAGLE v4.0 (https://faculty.washington.edu/browning/beagle/b4_0.html) with default settings and filtered once more for markers under a 5% MAF. Finally, a 6K (6.251.927) SNPs panel was defined for further analysis.

### Phenotypic data processing and statistics

On each panel, a random-effect analysis of variance was conducted on the whole population evaluated in both control and low N condition to test for genotype (G), treatment (T), and their interaction (GxT) effects with the following model using R/lme4:

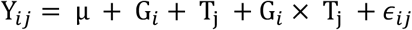

where Y_*ij*_ is the phenotype of genotype *i* in the treatment *j*; µis the overall mean, G_*i*_ is the genotypic effect of *i*^*th*^ genotype; T_*j*_is the effect of the *j*^*th*^ treatment; and ϵ_*ij*_ is the random residual effect with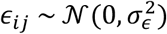.

Broad-sense heritability was calculated from the above model according to Cullis et al. (2006):

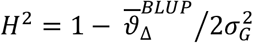

Where 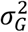 is the genotype variance, 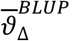 the average standard error of the genotypic BLUPs. Phenotypic plasticity traits were computed per accession as (y_*stress*_− y_*contorl*_)/ y_*contorl*._

This index was then used as a phenotype *per se* for QTL analysis. The average effect of the stress was reported as the mean relative variation and converted in percentage of increase or decrease due to the stress. The significance of the treatment effect was then calculated using a likelihood ratio test (R/*lmtest*) comparing the goodness-of-fit between two models, the first considering the treatment effect, the second one didn’tt taking it into account. A Box-Cox transformation (Box and Cox 1964) of the means calculated from replicates was applied to correct for heteroscedasticity and non-normality of error terms, before QTL mapping and GWAS analyses.

### Multi-parental QTL mapping

Linkage mapping in the MAGIC population was carried out with a set of 1,345 SNP markers selected from the genome resequencing of the eight parental lines (Pascual et al. 2015). The founder to offspring probabilities were predicted using the Hidden Markov algorithm implemented in calc_genoprob from R/qtl2 package (Broman et al. 2019) using the genetic map generated by Pascual et al. (2015). Then, the founder probabilities were used with the Haley–Knott regression model implemented in R/qtl2 for QTLs detection. Significance threshold was set to a LOD threshold of –log10(α/number of SNPs), where α was fixed at the 1% level. Then 2-LOD drop confidence intervals were calculated using the find_peaks() expanding it to a true marker on both sides of the QTL. Finally, candidate causal variants in accordance with the estimated founder effects were filtered as described in Pascual et al. (2015), by selecting the most contrasted pair of genotypes. For table summaries and transcriptome comparison, MAGIC variant-level *p*-values were grouped to obtain gene-level *p*-values by assuming a linear interpolation of the *p-values*.

### GWAS analysis

A univariate GWAS was performed by implementing the following linear mixed model in R/GENESYS (Gogarten et al. 2019) :

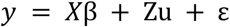

where y is the vector of phenotypic means for one environment, X is the molecular marker score matrix, β is the vector of marker effects, Z is an incidence matrix, u is the vector of random background polygenic effects with variance 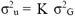 (where K is the kinship matrix, and 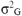 is the genetic variance), and ε is the vector of residuals.

The null model was fit using the NullModel() function using only the fixed-effect covariates. We included the first three eigenvectors estimated from the PCA based overall genotypic matrix using PLINK 2.0 (Purcell et al. 2007). Single-variant association tests were performed with assocTestSingle() function using Average Information REML (AIREML) procedure to estimate the variance components and score statistics. A LOD score of 5 was used as threshold. To estimate the proportion of variance explained (PVE) by lead SNP, we used the following formula proposed by Shim et al. (2015) using outputs from R/GENESYS:

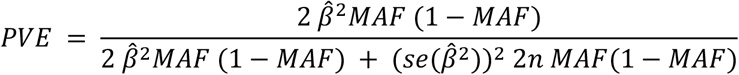

where 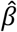 is the effect size, 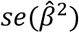 is the standard error for the effect size, *MAF* the minor allele frequency and *n* the sample size.

The GWAS-significant SNPs were binned into peaks based on linkage disequilibrium (LD), for QTL summary. In brief, for each trait/chromosome combination, the LD between the most significant SNP and all the others is computed. All the SNPs with a r^2^ > 0.5 are grouped together. The procedure was repeated for the remaining SNPs.

GWAS variant-level *p*-values were aggregated to obtain gene-level *p*-value, keeping the minimum p-value in a 10 kilobase window around the transcription start and stop sites, for candidate gene summaries and transcriptome comparison. Likewise, genes located within 10 kb of the flanking region of each significant SNP of a QTL were reported as candidate genes. Leaf NBI was excluded from GWAS analysis due to insufficient phenotypic records.

### RNA extraction

Four parental lines of the MAGIC population, Cervil, Ferum TMV, Levovil and LA1420 were selected for transcriptome analysis. For each accession, twelve plants were grown following the protocol described Abro et al. (2013), from February to June 2021 in a heated greenhouse at Avignon INRAE Centre (Avignon, France). Out of the twelve plants, six were fertigated with a solution containing 2 mM NO_3_^-^ (stress condition) and six with a solution containing 10 mM NO_3_^-^ (control conditions). The six plants were divided into three biological replicates of two plants. We performed RNA extraction on the three biological replicates obtained from pools of either young leaves sampled after the emergence of the fifth truss or tomato pericarp of at least five fruits picked at the turning stage. Total RNA was extracted, their purity and quality were assessed following the protocol described in Bineau et al. (2022).

### RNA sequencing, data processing and analysis

Library construction and sequencing (100 bp paired-end) strand of the 48 samples (4 accessions x 3 replicates x 2 organs x 2 treatments) were subcontracted to BGI genomics. The minimal, maximal, and average amounts of raw sequencing data per sample were estimated to be 20,389,318 bp, 21,047,855 bp and 20,493,477 bp, respectively. Raw data quality assessment, sequence cleaning, alignment and filtering were performed following the methodology described in Bineau et al. (2022). On average, 98.80 % of reads were properly paired (min = 95.36 %, max = 99.37 %). The insert size average was 493bp (min = 458bp, max = 559bp). The data from leaves and fruits were considered independently in the following analyses. We filtered out genes mapping on chromosome 0 and performed quality control, read count normalisation and sampling of genes expressed in the experiment using the workspace DiCoExpress with recommended parameters(Lambert et al. 2020). Twelve thousand eight hundreds twenty-seven (12,827) and 15,264 genes in fruit and leaf samples, respectively remained after the quality control (37.6 % and 44.8 % of all detected genes, respectively) for further analyses. DEG analysis was then performed on this subset using the workspace DiCoexpress with recommended parameters. After normalisation, to assess the differences of the transcriptome among the 4 lines and between N conditions, a principal component analysis (PCA) was performed. Then DEGs were detected for all possible contrasts - 29 - between the four lines and two N conditions. The p-values were corrected for multiple comparisons using the false discovery rate (Benjamini and Hochberg 1995) using a global threshold of 0.05. Gene enrichment was performed thanks to an adapted version of the Enrichment function of the DICoExpress workspace using the MAPMAN classification as reference (Provart and Zhu 2003).

## Data availability

The new genome reads corresponding to the 100 new accession sequences can be found in NCBI (https://www.ncbi.nlm.nih.gov/sra) database under accession number PRJNA1014227. RNA raw reads data can be accessed under number PRJNA817375.

The VCF file, RNAseq data and scripts used in the analysis are available at https://doi.org/10.57745/ELIUJ2.

All other data supporting the findings of this study are available in the supplementary data published online.

## Acknowledgement

We are thankful to Mélanie Andrin for taking care of the plants, to the vegetable resources centre (CRBLeg) of GAFL for keeping the seeds of the CC panel available and to the greenhouse staff of UE A2M “Arboriculture et Maraichage Méditerranéens”.

## Supplementary Materials

The following supporting information can be downloaded :

**Supplementary Table S1**: List of the 143 accessions of the diversity panel and their genome sequence references.

**Supplementary Table S2**: Phenotypic mean values measured for the MAGIC panel under control (C) and stress (S) treatment.

**Supplementary Table S3:** Phenotypic mean values measured for the diversity panel (core-collection) under control (C) and stress (S) treatment.

**Supplementary Table S4:** Summary of QTLs detected for fruit quality and agronomic traits in the the MAGIC panel

**Supplementary Table S5:** Summary of QTLs detected for fruit quality and agronomic traits in the diversity panel (GWAS)

**Supplementary Table S6:** List of genes used for MAGIC intersection analysis

**Supplementary Table S7:** List of genes used for GWAS intersection analysis

**Supplementary Table S8:** List of genes related to N metabolism and response to N stress, used for targeted differential expression analysis

**Supplementary figure S1**. Pictures of whole plants of the MAGIC parental lines under control (top) and stress (bottom) conditions at 70 days.

**Supplementary figure S2**. QTL profiles of N-related traits in the MAGIC population

**Supplementary figure S3**. Manhattan plots of N-related traits in the GWAS panel

**Supplementary figure S4**. Principal component analysis (PCA) of transcriptome-wide raw and normalized gene expression counts for leaves samples.

**Supplementary figure S5**. Principal component analysis (PCA) of transcriptome-wide raw and normalized gene expression counts for fruits samples.

**Supplementary figure S6**. Number of differentially expressed genes by contrast and by organ

**Supplementary figure S7**. Comparisons of the number of differentially expressed genes per organ between the Renau-Morata et al. (2021) study and the current study.

## References

Abenavoli MR, Longo C, Lupini A, Miller AJ, Araniti F, Mercati F, Princi MP, Sunseri F (2016) Phenotyping two tomato genotypes with different nitrogen use efficiency. Plant Physiol Biochem PPB 107:21–32.

Abro MA, Lecompte F, Bryone F, Nicot PC (2013) Nitrogen fertilization of the host plant influences production and pathogenicity of Botrytis cinerea secondary inoculum. Phytopathology 103:261–267.

Aci MM, Lupini A, Mauceri A, Sunseri F, Abenavoli MR (2021) New insights into N-utilization efficiency in tomato (Solanum lycopersicum L.) under N limiting condition. Plant Physiol Biochem 166:634–644.

Ahmad N, Su B, Ibrahim S, Kuang L, Tian Z, Wang X, Wang H, Dun X (2022) Deciphering the Genetic Basis of Root and Biomass Traits in Rapeseed (Brassica napus L.) through the Integration of GWAS and RNA-Seq under Nitrogen Stress. Int J Mol Sci 23:7958.

Albert E, Duboscq R, Latreille M, Santoni S, Beukers M, Bouchet J-P, Bitton F, Gricourt J, Poncet C, Gautier V, Jiménez-Gómez JM, Rigaill G, Causse M (2018) Allele-specific expression and genetic determinants of transcriptomic variations in response to mild water deficit in tomato. Plant J 96:635–650.

Albert E, Segura V, Gricourt J, Bonnefoi J, Derivot L, Causse M (2016) Association mapping reveals the genetic architecture of tomato response to water deficit: focus on major fruit quality traits. J Exp Bot 67:6413.

Asins MJ, Albacete A, Martinez-Andujar C, Pérez-Alfocea F, Dodd IC, Carbonell EA, Dieleman JA (2017) Genetic analysis of rootstock-mediated nitrogen (N) uptake and root-to-shoot signalling at contrasting N availabilities in tomato. Plant Sci 263:94–106.

Balazadeh S, Schildhauer J, Araújo WL, Munné-Bosch S, Fernie AR, Proost S, Humbeck K, Mueller-Roeber B (2014) Reversal of senescence by N resupply to N-starved Arabidopsis thaliana: transcriptomic and metabolomic consequences. J Exp Bot 65:3975–3992.

Bénard C, Gautier H, Bourgaud F, Grasselly D, Navez B, Caris-Veyrat C, Weiss M, Génard M (2009) Effects of low nitrogen supply on tomato (Solanum lycopersicum) fruit yield and quality with special emphasis on sugars, acids, ascorbate, carotenoids, and phenolic compounds. J Agric Food Chem 57:4112–4123.

Benjamini Y, Hochberg Y (1995) Controlling the False Discovery Rate: A Practical and Powerful Approach to Multiple Testing. J R Stat Soc Ser B Methodol 57:289–300.

Bineau E, Diouf I, Carretero Y, Duboscq R, Bitton F, Djari A, Zouine M, Causse M (2021) Genetic diversity of tomato response to heat stress at the QTL and transcriptome levels. Plant J 107:1213–1227.

Bineau E, Rambla J, Duboscq R, Corre M-N, Bitton F, Lugan R, Granell A, Plissonneau C, Causse M (2022) Inheritance of Secondary Metabolites and Gene Expression Related to Tomato Fruit Quality. Int J Mol Sci 23:6163.

Box GEP, Cox DR (1964) An Analysis of Transformations. J R Stat Soc Ser B Methodol 26:211–243.

Broman KW, Gatti DM, Simecek P, Furlotte NA, Prins P, SenŚ, Yandell BS, Churchill GA (2019) R/qtl2: Software for Mapping Quantitative Trait Loci with High-Dimensional Data and Multiparent Populations. Genetics 211:495–502.

Cao J, Zheng X, Xie D, Zhou H, Shao S, Zhou J (2022) Autophagic pathway contributes to low-nitrogen tolerance by optimizing nitrogen uptake and utilization in tomato. Hortic Res 9:uhac068.

Capua Y, Eshed Y (2017) Coordination of auxin-triggered leaf initiation by tomato LEAFLESS. Proc Natl Acad Sci 114:3246–3251.

Causse M, Desplat N, Pascual L, Le Paslier M-C, Sauvage C, Bauchet G, Bérard A, Bounon R, Tchoumakov M, Brunel D, Bouchet J-P (2013) Whole genome resequencing in tomato reveals variation associated with introgression and breeding events. BMC Genomics 14. https://www.ncbi.nlm.nih.gov/pmc/articles/PMC4046683/ (30 August 2019, date last accessed).

Chen S, Zhou Y, Chen Y, Gu J (2018) fastp: an ultra-fast all-in-one FASTQ preprocessor. Bioinformatics 34:i884–i890.

Cullis BR, Smith AB, Coombes NE (2006) On the design of early generation variety trials with correlated data. J Agric Biol Environ Stat 11:381.

Curci PL, Aiese Cigliano R, Zuluaga DL, Janni M, Sanseverino W, Sonnante G (2017) Transcriptomic response of durum wheat to nitrogen starvation. Sci Rep 7:1176.

de Souza R, Grasso, Peña-Fleitas MT, Gallardo, Thompson RB, Padilla F (2020) Effect of Cultivar on Chlorophyll Meter and Canopy Reflectance Measurements in Cucumber. Sensors 20:509.

FAOSTAT. https://www.fao.org/faostat/en/#data/QCL (23 October 2023, date last accessed).

Fu Y, Yi H, Bao J, Gong J (2015) LeNRT2.3 functions in nitrate acquisition and long-distance transport in tomato. FEBS Lett 589:1072–1079.

Gogarten SM, Sofer T, Chen H, Yu C, Brody JA, Thornton TA, Rice KM, Conomos MP (2019) Genetic association testing using the GENESIS R/Bioconductor package. Bioinforma Oxf Engl 35:5346–5348.

Guo M, Yang F, Liu C, Zou J, Qi Z, Fotopoulos V, Lu G, Yu J, Zhou J (2022) A single-nucleotide polymorphism in WRKY33 promoter is associated with the cold sensitivity in cultivated tomato. New Phytol 236:989–1005.

Incrocci L, Thompson RB, Fernandez-Fernandez MD, De Pascale S, Pardossi A, Stanghellini C, Rouphael Y, Gallardo M (2020) Irrigation management of European greenhouse vegetable crops. Agric Water Manag 242:106393.

Islam MdN, Hasanuzzaman ATM, Zhang Z-F, Zhang Y, Liu T-X (2017) High Level of Nitrogen Makes Tomato Plants Releasing Less Volatiles and Attracting More Bemisia tabaci (Hemiptera: Aleyrodidae). Front Plant Sci 8:466.

Johansen K, Morton MJL, Malbeteau Y, Aragon B, Al-Mashharawi S, Ziliani MG, Angel Y, Fiene G, Negrão S, Mousa MAA, Tester MA, McCabe MF (2020) Predicting Biomass and Yield in a Tomato Phenotyping Experiment Using UAV Imagery and Random Forest. Front Artif Intell 3. https://www.frontiersin.org/articles/10.3389/frai.2020.00028 (14 December 2022, date last accessed).

Khan M, Struik P, Putten P, Yin X, Eck HJ, Eeuwijk F (2010) Genetic variation in potato (Solanum tuberosum L.) canopy development: A model approach using standard cultivars and a segregating population. Eur Potato J 53:199 –252.

Köster J, Rahmann S (2012) Snakemake--a scalable bioinformatics workflow engine. Bioinforma Oxf Engl 28:2520–2522.

Krapp A, Berthomé R, Orsel M, Mercey-Boutet S, Yu A, Castaings L, Elftieh S, Major H, Renou J-P, Daniel-Vedele F (2011) Arabidopsis roots and shoots show distinct temporal adaptation patterns toward nitrogen starvation. Plant Physiol 157:1255–1282.

Kurtzer GM, Sochat V, Bauer MW (2017) Singularity: Scientific containers for mobility of compute. PloS One 12:e0177459.

Lambert I, Paysant-Le Roux C, Colella S, Martin-Magniette M-L (2020) DiCoExpress: a tool to process multifactorial RNAseq experiments from quality controls to co-expression analysis through differential analysis based on contrasts inside GLM models. Plant Methods 16:68.

Larbat R, Paris C, Le Bot J, Adamowicz S (2014) Phenolic characterization and variability in leaves, stems and roots of Micro-Tom and patio tomatoes, in response to nitrogen limitation. Plant Sci 224:62–73.

Li H, Handsaker B, Wysoker A, Fennell T, Ruan J, Homer N, Marth G, Abecasis G, Durbin R, 1000 Genome Project Data Processing Subgroup (2009) The Sequence Alignment/Map format and SAMtools. Bioinformatics 25:2078–2079.

Liu Q, Wu K, Song W, Zhong N, Wu Y, Fu X (2022) Improving Crop Nitrogen Use Efficiency Toward Sustainable Green Revolution. Annu Rev Plant Biol 73:523–551.

Luo L, Zhang Y, Xu G (2020) How does nitrogen shape plant architecture? J Exp Bot 71:4415–4427.

Magán JJ, Gallardo M, Thompson RB, Lorenzo P (2008) Effects of salinity on fruit yield and quality of tomato grown in soil-less culture in greenhouses in Mediterranean climatic conditions. Agric Water Manag 95:1041–1055.

Méndez-Cifuentes A, Valdez-Aguilar LA, Cadena-Zapata M, González-Fuentes JA, Hernández-Maruri JA, Alvarado-Camarillo D (2020) Water and Fertilizer Use Efficiency in Subirrigated Containerized Tomato. Water 12:1313.

Minotti PL, Halseth DE, Sieczka JB (1994) Field Chlorophyll Measurements to Assess the Nitrogen Status of Potato Varieties. HortScience 29:1497–1500.

Moll RH, Kamprath EJ, Jackson WA (1982) Analysis and Interpretation of Factors Which Contribute to Efficiency of Nitrogen Utilization1. Agron J 74:562–564.

Monostori I, Árendás T, Hoffman B, Galiba G, Gierczik K, Szira F, Vágújfalvi A (2016) Relationship between SPAD value and grain yield can be affected by cultivar, environment and soil nitrogen content in wheat. Euphytica 211:103–112.

Noguchi K (2006) Effects of Light Intensity and Carbohydrate Status on Leaf and Root Respiration. In: Plant Respiration. pp 63–83.

Pascual L, Albert E, Sauvage C, Duangjit J, Bouchet J-P, Bitton F, Desplat N, Brunel D, Le Paslier M-C, Ranc N, Bruguier L, Chauchard B, Verschave P, Causse M (2016) Dissecting quantitative trait variation in the resequencing era: complementarity of bi-parental, multi-parental and association panels. Plant Sci 242:120–130.

Pascual L, Desplat N, Huang BE, Desgroux A, Bruguier L, Bouchet J-P, Le QH, Chauchard B, Verschave P, Causse M (2015) Potential of a tomato MAGIC population to decipher the genetic control of quantitative traits and detect causal variants in the resequencing era. Plant Biotechnol J 13:565–577.

Peña-Fleitas MT, Gallardo M, Padilla FM, Rodríguez A, Thompson RB (2021) Use of a Portable Rapid Analysis System to Measure Nitrate Concentration of Nutrient and Soil Solution, and Plant Sap in Greenhouse Vegetable Production. Agronomy 11:819.

Provart N, Zhu T (2003) A Browser-based Functional Classification SuperViewer for Arabidopsis Genomics. Curr Comput Mol Biol 2003

Purcell S, Neale B, Todd-Brown K, Thomas L, Ferreira MAR, Bender D, Maller J, Sklar P, de Bakker PIW, Daly MJ, Sham PC (2007) PLINK: a tool set for whole-genome association and population-based linkage analyses. Am J Hum Genet 81:559–575.

Ren T, Li Y, Miao T, Hassan W, Zhang J, Wan Y, Cai A (2022) Characteristics and Driving Factors of Nitrogen-Use Efficiency in Chinese Greenhouse Tomato Cultivation. Sustainability 14:805.

Renau-Morata B, Cebolla-Cornejo J, Carrillo L, Gil-Villar D, Martí R, Jiménez-Gómez JM, Granell A, Monforte AJ, Medina J, Molina RV, Nebauer SG (2024) Identification of Solanum pimpinellifolium genome regions for increased resilience to nitrogen deficiency in cultivated tomato. Sci Hortic 323:112497.

Renau-Morata B, Molina R-V, Minguet EG, Cebolla-Cornejo J, Carrillo L, Martí R, García-Carpintero V, Jiménez-Benavente E, Yang L, Cañizares J, Canales J, Medina J, Nebauer SG (2021) Integrative Transcriptomic and Metabolomic Analysis at Organ Scale Reveals Gene Modules Involved in the Responses to Suboptimal Nitrogen Supply in Tomato. Agronomy 11:1320.

Rosa-Martínez E, Adalid AM, Alvarado LE, Burguet R, García-Martínez MD, Pereira-Dias L, Casanova C, Soler E, Figàs MR, Plazas M, Prohens J, Soler S (2021) Variation for Composition and Quality in a Collection of the Resilient Mediterranean ‘de penjar’ Long Shelf-Life Tomato Under High and Low N Fertilization Levels. Front Plant Sci 12. https://www.frontiersin.org/articles/10.3389/fpls.2021.633957 (3 October 2023, date last accessed).

Shim H, Chasman DI, Smith JD, Mora S, Ridker PM, Nickerson DA, Krauss RM, Stephens M (2015) A Multivariate Genome-Wide Association Analysis of 10 LDL Subfractions, and Their Response to Statin Treatment, in 1868 Caucasians. PLoS ONE 10:e0120758.

Shin S-Y, Jeong JS, Lim JY, Kim T, Park JH, Kim J-K, Shin C (2018) Transcriptomic analyses of rice (Oryza sativa) genes and non-coding RNAs under nitrogen starvation using multiple omics technologies. BMC Genomics 19:532.

Snyder CS, Bruulsema TW, Jensen TL, Fixen PE (2009) Review of greenhouse gas emissions from crop production systems and fertilizer management effects. Agric Ecosyst Environ 133:247–266.

Sung J, Lee S, Lee Y, Ha S, Song B, Kim T, Waters BM, Krishnan HB (2015) Metabolomic profiling from leaves and roots of tomato (Solanum lycopersicum L.) plants grown under nitrogen, phosphorus or potassium-deficient condition. Plant Sci Int J Exp Plant Biol 241:55–64.

Sunseri F, Aci MM, Mauceri A, Caldiero C, Puccio G, Mercati F, Abenavoli MR (2023) Short-term transcriptomic analysis at organ scale reveals candidate genes involved in low N responses in NUE-contrasting tomato genotypes. Front Plant Sci 14. https://www.frontiersin.org/articles/10.3389/fpls.2023.1125378 (23 October 2023, date last accessed).

The SV, Snyder R, Tegeder M (2021) Targeting Nitrogen Metabolism and Transport Processes to Improve Plant Nitrogen Use Efficiency. Front Plant Sci 11:628366.

Truffault V, Ristorto M, Brajeul E, Vercambre G, Gautier H (2019) To Stop Nitrogen Overdose in Soilless Tomato Crop: A Way to Promote Fruit Quality without Affecting Fruit Yield. Agronomy 9:80.

Urbanczyk-Wochniak E, Fernie AR (2005) Metabolic profiling reveals altered nitrogen nutrient regimes have diverse effects on the metabolism of hydroponically-grown tomato (Solanum lycopersicum) plants. J Exp Bot 56:309–321.

Vallarino JG, Shameer S, Zhang Y, Ratcliffe RG, Sweetlove LJ, Fernie AR (2022) Leaf specific overexpression of a mitochondrially-targeted glutamine synthetase in tomato increased assimilate export resulting in earlier fruiting and elevated yield. bioRxiv:2022.06.28.497938.

Wang X, Zou C, Gao X, Guan X, Zhang W, Zhang Y, Shi X, Chen X (2018) Nitrous oxide emissions in Chinese vegetable systems: A meta-analysis. Environ Pollut 239:375–383.

Xu G, Fan X, Miller AJ (2012) Plant nitrogen assimilation and use efficiency. Annu Rev Plant Biol 63:153–182.

Xun Z, Guo X, Li Y, Wen X, Wang C, Wang Y (2020) Quantitative proteomics analysis of tomato growth inhibition by ammonium nitrogen. Plant Physiol Biochem 154:129–141.

Yifhar T, Pekker I, Peled D, Friedlander G, Pistunov A, Sabban M, Wachsman G, Alvarez JP, Amsellem Z, Eshed Y (2012) Failure of the Tomato Trans-Acting Short Interfering RNA Program to Regulate AUXIN RESPONSE FACTOR3 and ARF4 Underlies the Wiry Leaf Syndrome[C][W]. Plant Cell 24:3575–3589.

Zhang X, Ma Q, Li F, Ding Y, Yi Y, Zhu M, Ding J, Li C, Guo W, Zhu X (2021) Transcriptome Analysis Reveals Different Responsive Patterns to Nitrogen Deficiency in Two Wheat Near-Isogenic Lines Contrasting for Nitrogen Use Efficiency. Biology 10:1126.

Zhu G, Wang S, Huang Z, Zhang S, Liao Q, Zhang C, Lin T, Qin M, Peng M, Yang C, Cao X, Han X, Wang X, van der Knaap E, Zhang Z, Cui X, Klee H, Fernie AR, Luo J, Huang S (2018) Rewiring of the Fruit Metabolome in Tomato Breeding. Cell 172:249-261.e12.

