## Supplementary figures for "Integration of QTL and transcriptome approaches for the identification of genes involved in tomato response to nitrogen deficiency"

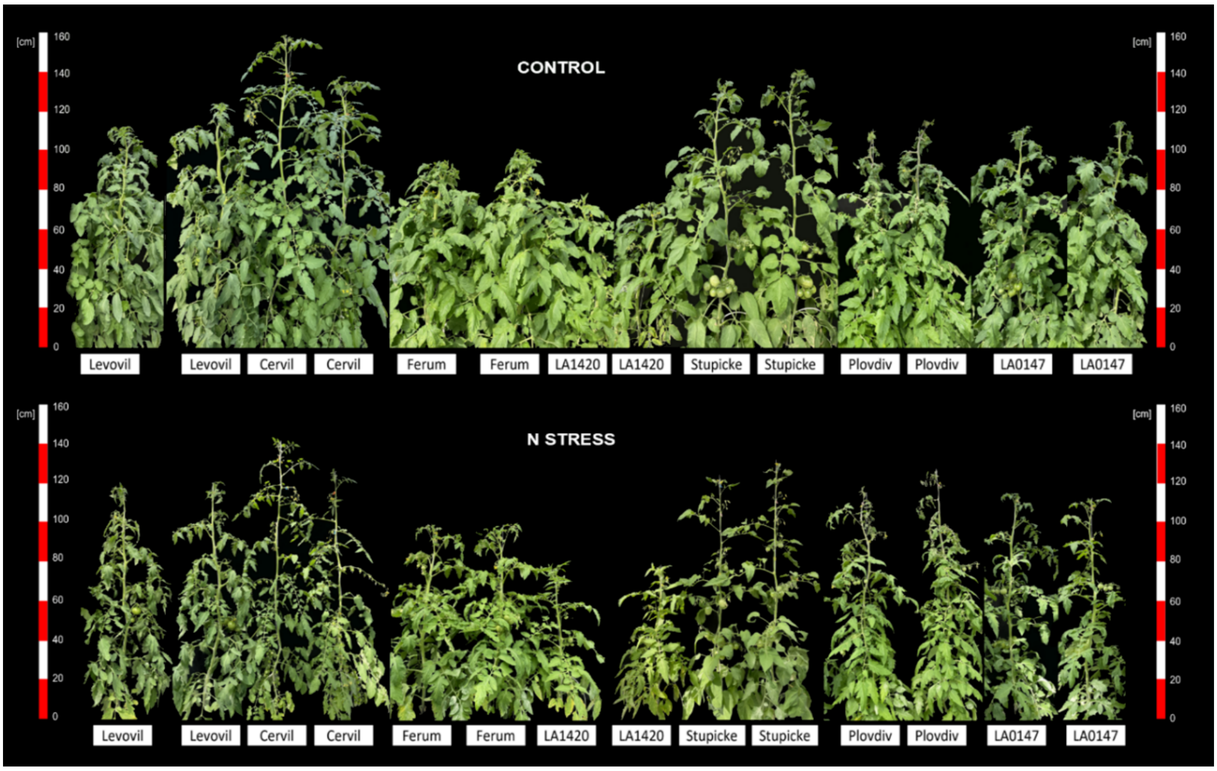


**Supplementary figure S1.** Pictures of whole plants of the MAGIC parental lines under control (top) and stress (bottom) conditions at 70 days.

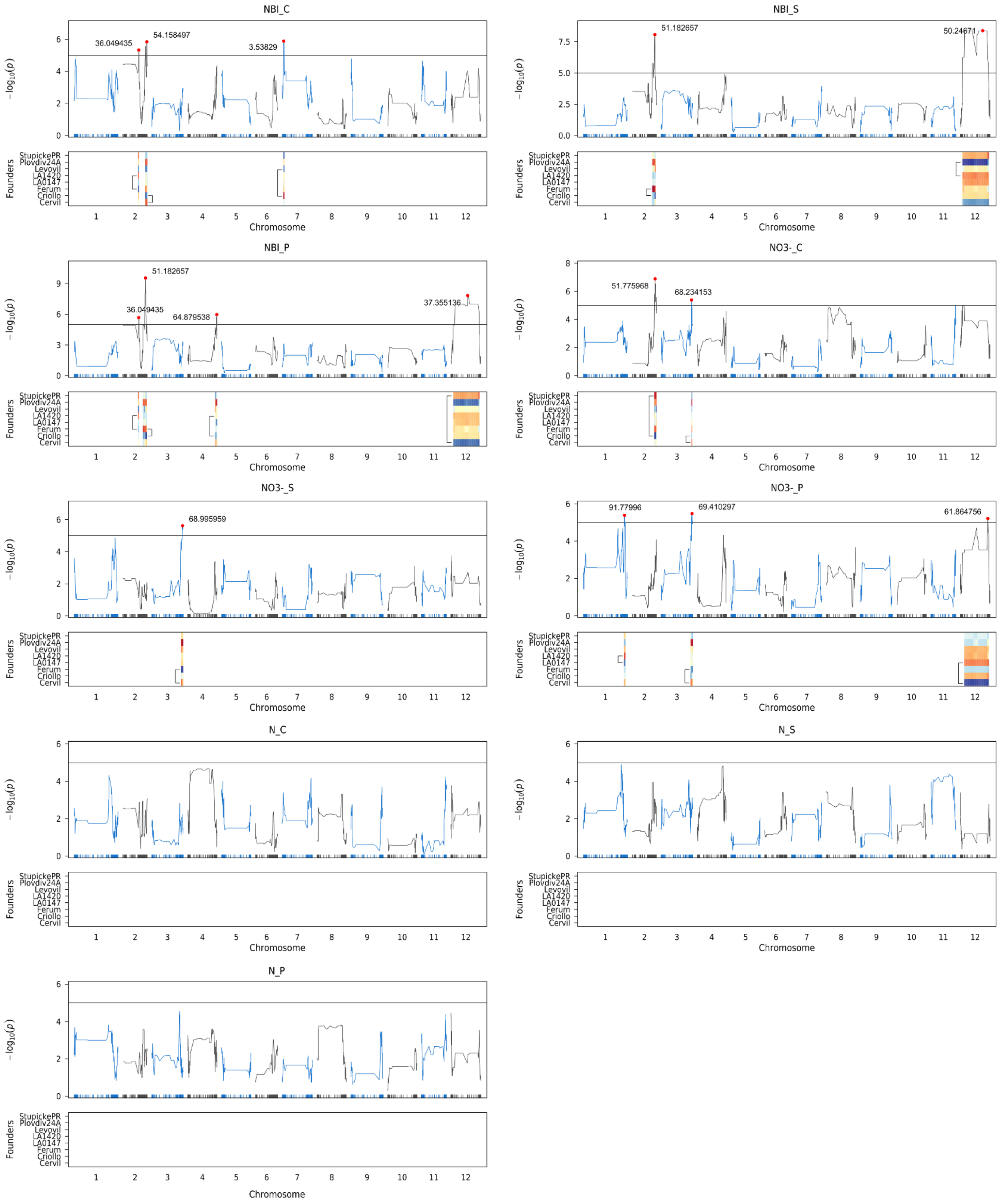


**Supplementary figure S2. QTL profiles of N-related traits in the MAGIC population.**
The x-axis represents physical distance according to ITAG4.0. The y-axis of the upper panel represents LOD scores. The lower panel corresponds to the estimated founder effects in the QTL confidence interval. Brackets denote the most contrasting pair of allelic effects used to filter candidate genes.

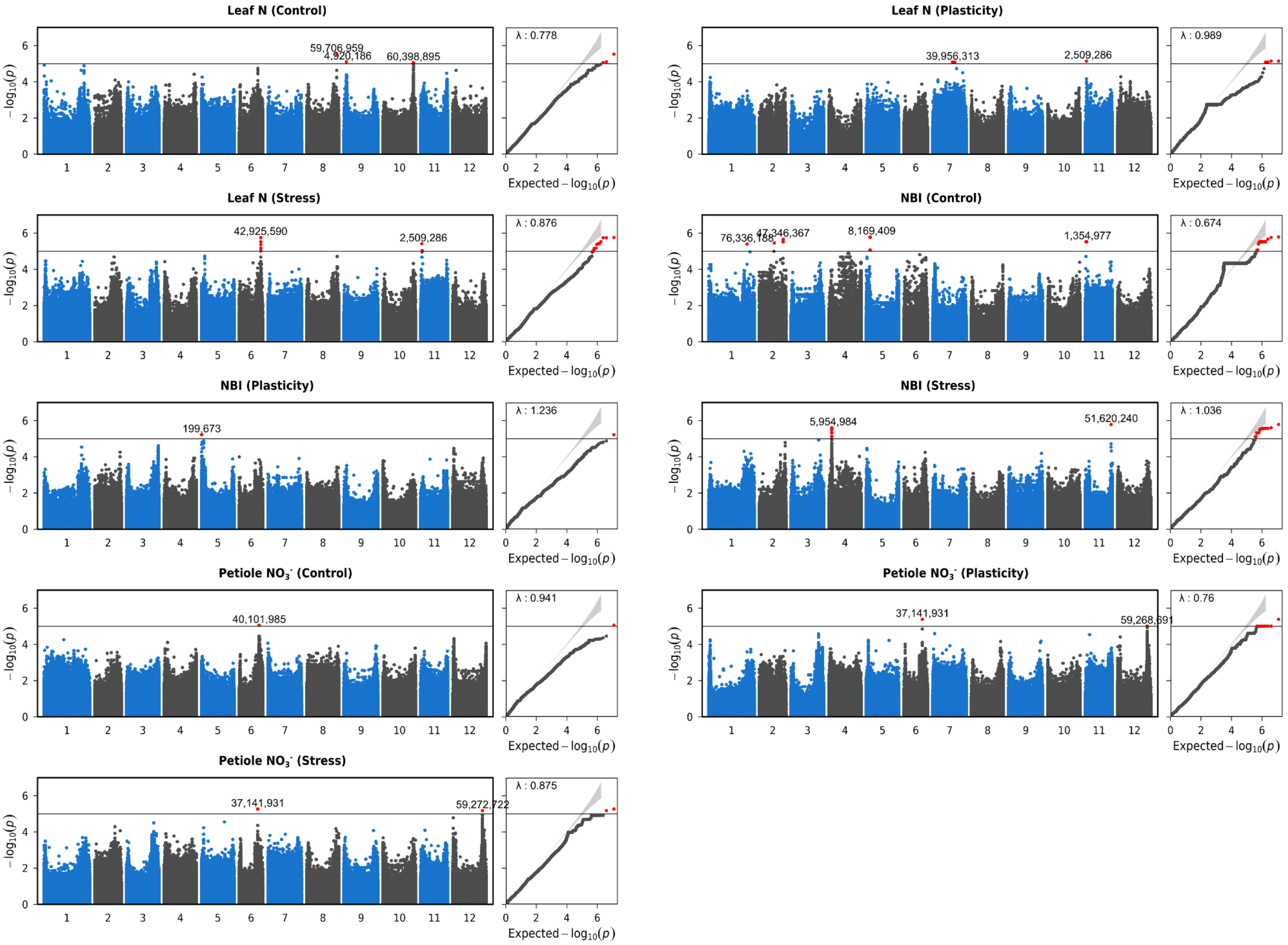


**Supplementary figure S3. Manhattan plots of N-related traits in the GWAS panel**
The x-axis represents physical distance according to ITAG4.0. The y-axis represents LOD scores. Red dots indicate QTLs with $-\log_{10}(pvalue) > 5$. Positions of lead SNP are indicated in bp.
.

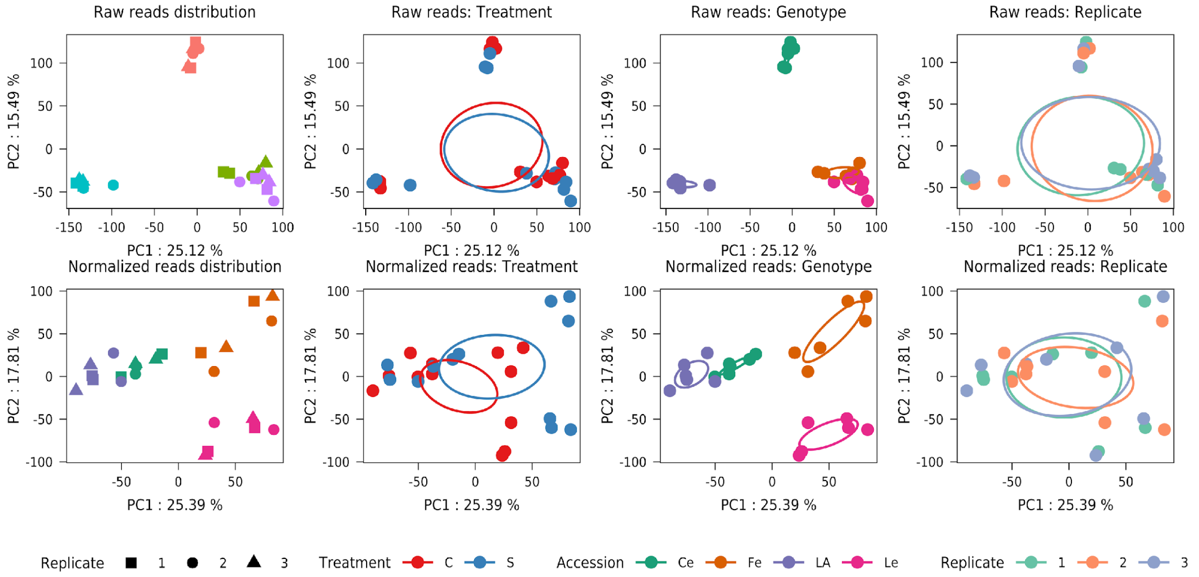


**Supplemental figure S4**. Principal component analysis (PCA) of transcriptome-wide raw and normalized gene expression counts for **leaves samples**.
**Ce** : Cervil, **Fe** : Ferum, **LA** : LA1420, **Le** : Levovil, **C** : Control condition, **S** : Stress condition.

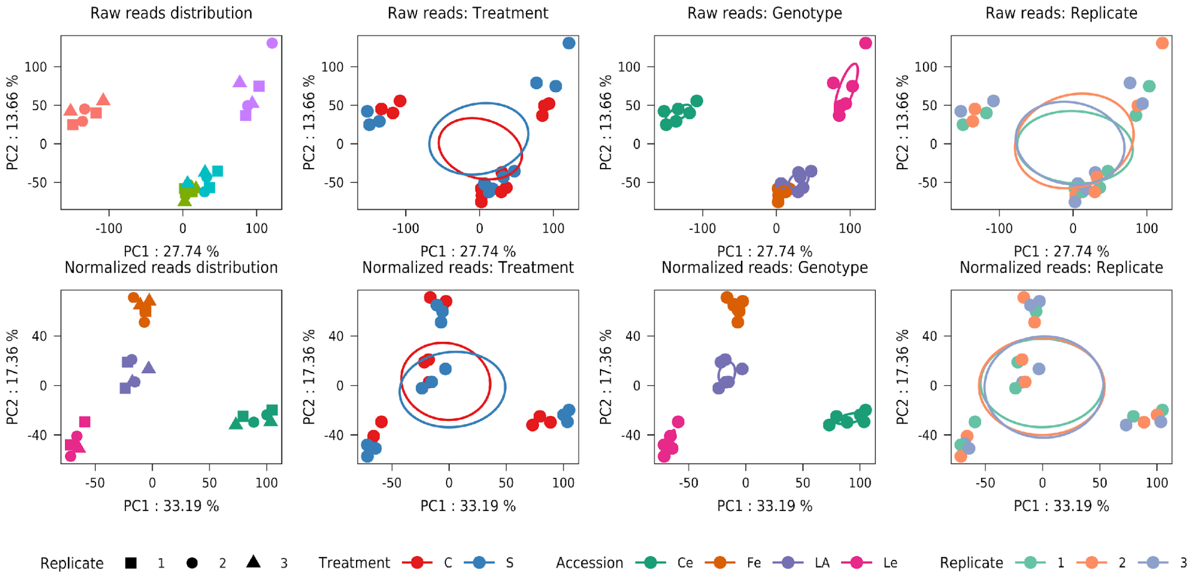


**Supplemental figure S5**. Principal component analysis (PCA) of transcriptome-wide raw and normalized gene expression counts for **fruit samples**.
**Ce** : Cervil, **Fe** : Ferum, **LA** : LA1420, **Le** : Levovil, **C** : Control condition, **S** : Stress condition.

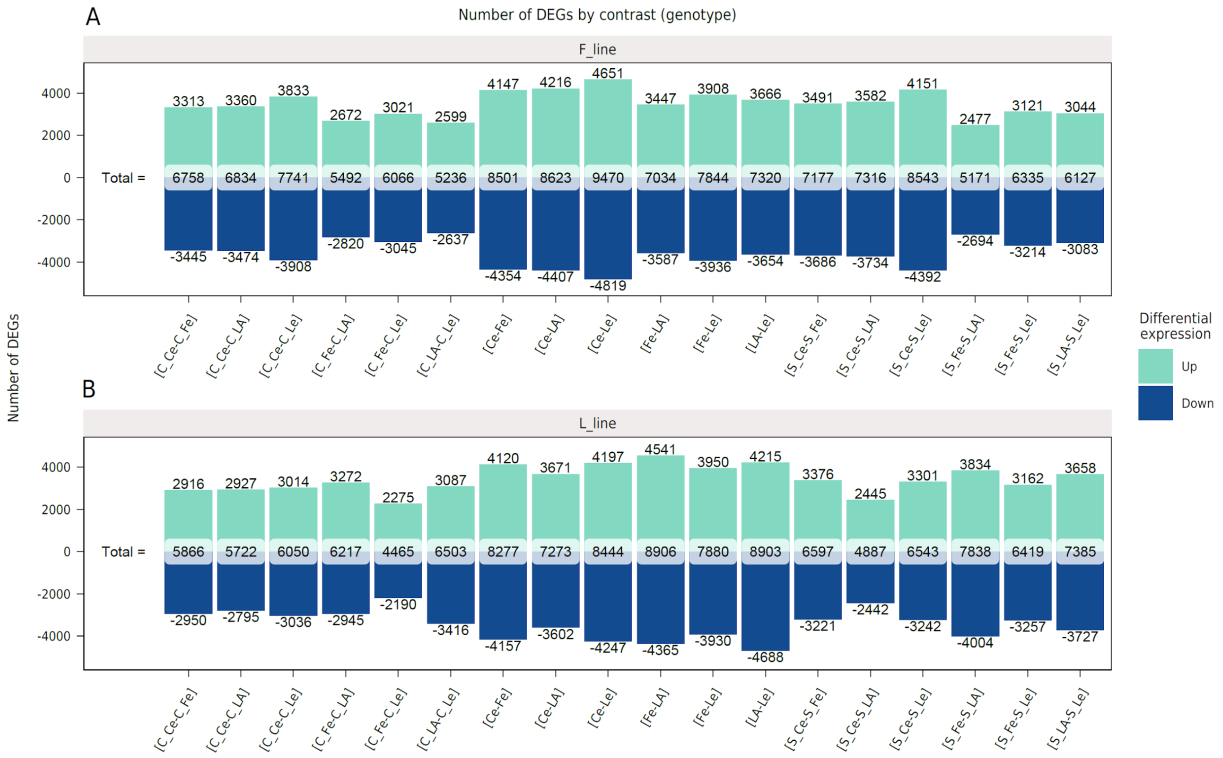


**Supplemental figure S6**. Number of differentially expressed genes by contrast and by organ (A. Fruits; B. Leaves). **Ce** : Cervil, **Fe** : Ferum, **LA** : LA1420, **Le** : Levovil, **C** : Control condition, **S** : Stress condition.

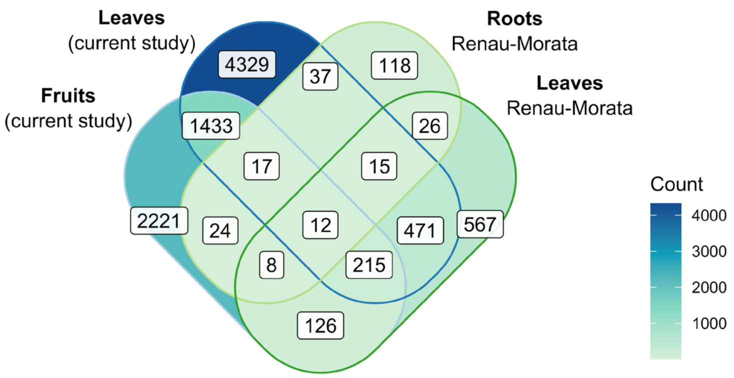


**Supplementary figure S7**. Comparisons of the number of differentially expressed genes per organ between the Renau-Morata study (2021) and the current study.
